# Spatial Co-transcriptomics Reveals Discrete Stages of the Arbuscular Mycorrhizal Symbiosis

**DOI:** 10.1101/2023.08.02.551648

**Authors:** Karen Serrano, Margaret Bezrutczyk, Danielle Goudeau, Thai Dao, Ronan O’Malley, Rex R. Malmstrom, Axel Visel, Henrik Scheller, Benjamin Cole

## Abstract

The symbiotic interaction of plants with arbuscular mycorrhizal fungi (AM fungi) is ancient and widespread. Plants provide AM fungi with carbon in exchange for nutrients and water, making this interaction a prime target for crop improvement. However, plant-fungal interactions are restricted to a small subset of root cells, precluding the application of most conventional functional genomic techniques to study the molecular bases of these interactions. Here we used single-nucleus and spatial RNA sequencing to explore both *M. truncatula* and *R. irregularis* transcriptomes in AM symbiosis at cellular and spatial resolution. Integrated spatially-registered single-cell maps of interacting cells revealed major infected and uninfected plant root cell types. We observed that cortex cells exhibit distinct transcriptome profiles during different stages of colonization by AM fungi, indicating dynamic interplay between both organisms during establishment of the cellular interface enabling successful symbiosis. Our study provides insight into a symbiotic relationship of major agricultural and environmental importance and demonstrates a paradigm combining single-cell and spatial transcriptomics for the analysis of complex organismal interactions.

## Introduction

Arbuscular mycorrhizal (AM) fungi occur in all major terrestrial ecosystems^1^. They are fundamental to agricultural production due to their ability to provide plants with a variety of nutrients, particularly non-renewable phosphorus, as well as resistance to abiotic stress^2^ and pathogens^3^. Plant hosts reward these services by transferring up to 20% of their photosynthate in the form of carbohydrates and lipids^4^ and enable the formation of an extensive extraradical mycelium in the soil.

To colonize their host, AM fungi extend intraradical hyphae between root epidermal cells deep into the cortex. AM fungi then penetrate cortex cells, where they form highly branched hyphal structures called arbuscules^5^. In response, the host plant restructures the cortex cell to accommodate and build a peri-arbuscular membrane around the arbuscule^6^. Nutrient and metabolite exchange between the two symbiotic partners occurs across the periarbuscular membrane via transport proteins localized specifically to this membrane, such as the essential phosphate transporter, *PHOSPHATE TRANSPORTER 4 (MtPT4)*^7^. The recruitment of AM fungi to the root surface and the development, maintenance, and senescence of arbuscules are tightly regulated processes that require intense coordination and communication between the two species.

Despite many years of research, the identity and function of many transcripts differentially expressed by the plant and fungus during this interaction are not yet characterized. Multiple characteristics of AM symbiosis complicate their study by traditional molecular genetic and transcriptomic approaches. As obligate biotrophs, AM fungi cannot complete their full life cycles in asymbiotic conditions, nor can they be readily cultured independently in the laboratory.

Additionally, mycorrhizal colonization occurs in a highly asynchronous manner, with many developmental stages existing simultaneously within the cortex, limiting the ability of whole-root transcriptome profiling to differentiate between gene expression associated with each stage^8^. Arbuscules are extremely transient structures, lasting only a few days before senescence^9^, confounding efforts to distinguish discrete phases of interaction. Furthermore, the collapse of arbuscules and appearance of vesicles or new spores indicates that the plant and fungus are assimilating the exchanged nutrients and continuing the symbiosis, which creates the need to analyze root cells that appear to be non-colonized just as important as those which contain arbuscules^10^. Several groups have elegantly addressed these challenges using laser capture microdissection (LCM) to obtain transcriptomes of cortex cells visually confirmed to be directly adjacent to fungal appressoria (early stage) and colonized cortex cells (late stage)^11, 12^. One disadvantage of LCM is that it limits the investigation to cell types already known to be involved in the interaction, which creates the need for an approach that will allow for unbiased investigation of all cell types within the root.

The recent and rapidly increasing adoption of single-cell RNA-sequencing (scRNA-seq) or single-nuclei RNA-sequencing (snRNA-seq), with their potential to identify novel cell types and states, model developmental trajectories, and analyze transcriptional activity in individual cells^13^, have revolutionized the field of plant biology. In both scRNA-seq and snRNA-seq, investigation of the transcriptomes from all cell types is possible, rather than requiring the investigator to manually select individual cells, as is the case with LCM. Moreover, snRNA-seq has the advantage over scRNA-seq in that its quick protocol is robust to diverse organisms and tissue types. In addition, certain cell types are preferentially released as protoplasts during enzymatic digestion for scRNA-seq, but nuclei are extracted more uniformly across cell types, leading to a more representative population of cell types in snRNA-seq datasets for plant tissues^14, 15^. In both scRNA-seq and snRNA-seq, the spatial context of gene expression is lost upon dissociation of cells or nuclei from the tissue. Spatial transcriptomics allows for sequencing of cell transcriptomes within the tissue context in an untargeted manner via the use of barcoded oligo (dT) arrays to capture polyadenylated transcripts, adding a novel dimension to the data^16^.

This study applied single-nuclei and spatial transcriptomics to the interaction between the model legume *Medicago truncatula* and the AM fungus *Rhizophagus irregularis* to create a 2-dimensional integrated map of plant and fungal transcriptomes during symbiosis. We provide an unbiased spatial and single-nuclei transcriptomics dataset that simultaneously profiled a multi-kingdom interaction. The spatially-resolved transcriptome provides insight into coordinated gene expression occurring between the two partners across all major *M. truncatula* root cell types. This transcriptomic map represents a novel resource for AM fungi research and demonstrates the value of novel multi-omics approaches in answering biological questions.

## Results

### Single-nuclei RNA sequencing identifies known M. truncatula root cell types

To gain a comprehensive transcriptional profile of the plant/AM fungal interaction, we performed snRNA-seq and spatial RNA-seq on *M. truncatula* roots which were mock-inoculated or inoculated with the AM fungus *R. irregularis.* We isolated and purified nuclei from *M. truncatula* roots by flow sorting (FANS, Fig. 1a) before loading the suspension onto a microfluidic chip for snRNA-seq profiling^17^. Of the seven datasets generated (Fig. S4, Table S1), we selected five high quality datasets for further characterization, resulting in a final dataset of 16,890 nuclei with an average of 1,120 mRNA molecules per cell after quality filtering. This dataset was then processed using Seurat^18, 19^, resulting in a two-dimensional Uniform Manifold Approximation and Projection plot (UMAP, a dimension reduction technique for visualization; Fig. 1b). Unsupervised clustering was performed to group cells into 16 clusters and cell identities were then assigned based on the expression of established marker genes for different cell types (Table S2) derived from *Arabidopsis thaliana* root single cell^20–24^ datasets, as well as a recently-published rhizobia-colonized *M. truncatula* root tip^25^ single-nuclei dataset.

**Figure 1:**
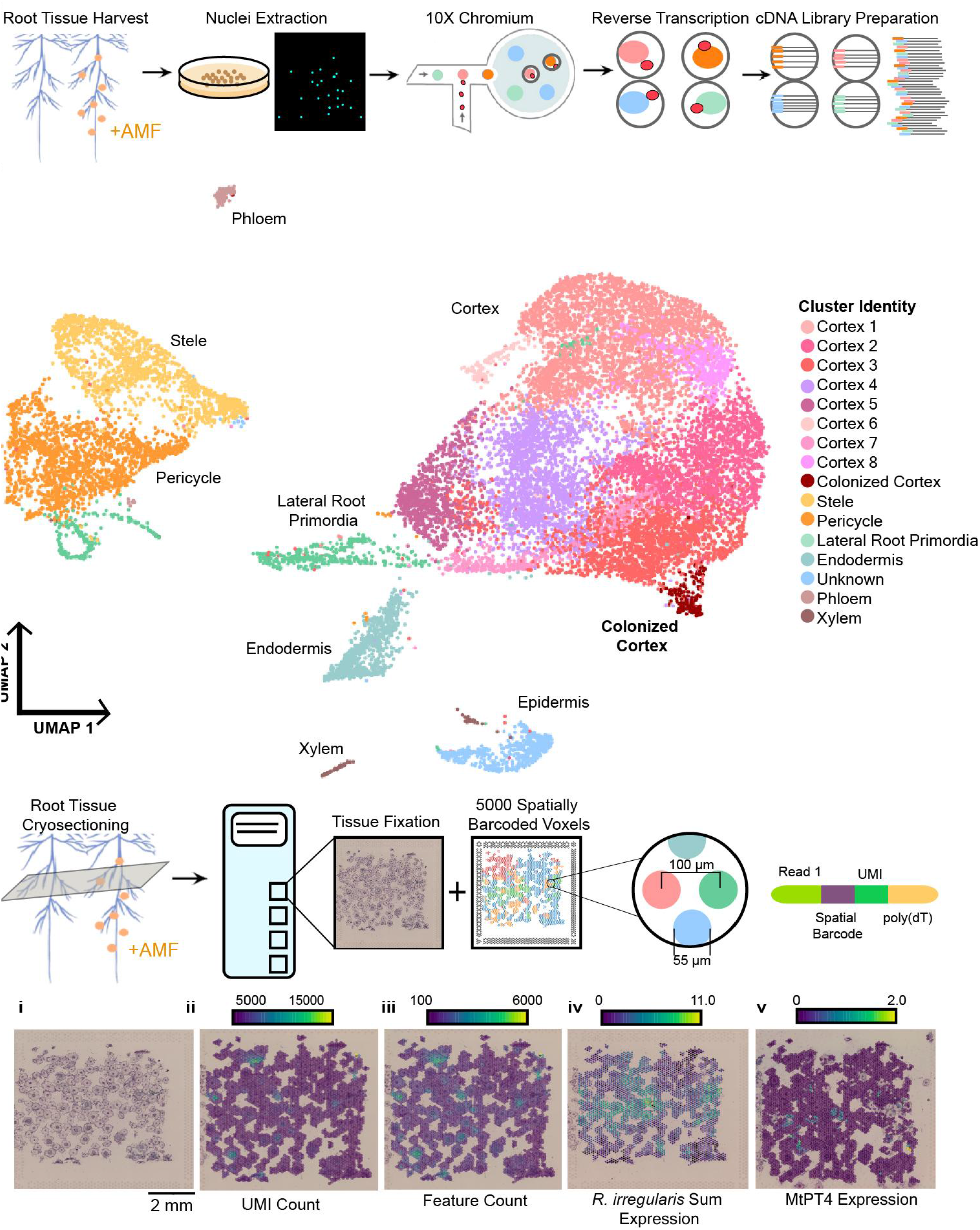
Single nuclei and spatial RNA profiling in *M. truncatula* roots colonized by *R. irregularis.* **a,** Overview of approach. *M. truncatula* root tissue is flash frozen for nuclei extraction and subsequent single-nuclei RNA sequencing using the 10X Genomics Chromium platform. Intact single nuclei are emulsified with gel beads containing barcoded oligonucleotides within a microfluidic chamber, resulting in a barcoded cDNA library after reverse transcription. **b,** UMAP coordinates of 16890 *M. truncatula* nuclei from three AM-colonized root harvests timepoints clustered by similarity in transcriptional profiles. The identities of 16 unique clusters are represented by different colors. **c,** *M. truncatula* root tissue is flash frozen to create 16 µm thick cryosections, each containing numerous root cross-sections. Cryosections are fixed to capture areas, each of which is equipped with ∼5000 spatially-barcoded voxels at a resolution of 55 µm. **d,** A single capture area of root cross sections from *M. truncatula* infected with *R. irregularis* at 28 dpi. i, Image of root cross sections within capture area. UMI count (ii) and feature count (iii) overlaid onto spots underlying tissue. iv, Expression pattern of all *R. irregularis* transcripts captured (scale = log2 of UMI counts). v, Expression pattern of the arbuscule marker gene post-imputation, *MtPT4* (*PHOSPHATE TRANSPORTER 4*), exhibiting overlap in spots with the highest expression of fungal transcripts (scale = log2(UMI counts)). Visualization done in Loupe Browser.

Cell identities of the 16 clusters were determined using a combination of known Medicago marker genes, as well as homologs of Arabidopsis marker genes (Text S1, Table S2, Fig. S3). Importantly, nine clusters were found to represent cortex cells, with one additional cluster composed of cortex cells that were colonized by *R. irregularis*. We observed a large number of cortex cells (11,298), and a much smaller number (174) of colonized cortex cells, indicating that we not only broadly recapitulated the cell type distribution of this tissue type, but also were able to estimate the fraction of colonized cells. All major medicago root cell types were found in our datasets, including vascular, endodermal, and meristematic cells (Fig. 1b, Text S1, Table S2, Fig. S3)

### Spatial transcriptomic profiling simultaneously captures plant and fungal transcripts

In order to investigate gene expression from both symbiotic partners simultaneously, we performed spatial transcriptomic profiling on inoculated and mock-inoculated *M. truncatula* roots at 28 dpi (Fig. 1c). An example capture area from an inoculated plant is shown in Fig. 1d panel I, which displays numerous lateral root cross sections fixed and stained on the capture area prior to tissue permeabilization. On average, inoculated capture areas resulted in 20,333 and 5,084 features mapping to the *M. truncatula* and *R. irregularis* genomes respectively, while mock-inoculated capture areas resulted in an average of 21,987 (*M. truncatula*) and 23 (*R. irregularis*) features. Spatial UMI (Fig. 4d, panel II) and feature (Fig. 4d, panel III) distributions indicated a uniform capture of transcripts across the expected areas of each cryosection, with few hotspots of fungal colonization associated with increased transcript counts from *R.irregularis* (Fig.1d, panel IV) and increased expression of *MtPT4*^9^ (Fig.1d, panel V). The spatial transcriptomics technology relies on polyadenylated transcript capture via oligo(dT) primer sequences which allowed the unbiased capture of *R. irregularis* and *M. truncatula* transcripts simultaneously. Analyzing the spatial distribution of all *R. irregularis* transcript expression within each of the nine inoculated capture areas (Fig. 2a, panel I) and comparing it to that of all *M. truncatula* transcripts (Fig 2a, panels II and III), distinct patterns of mRNA capture and expression between the two species are evident, indicating the successful spatially resolved capture of transcriptomes during symbiotic interaction.

**Figure 2.**
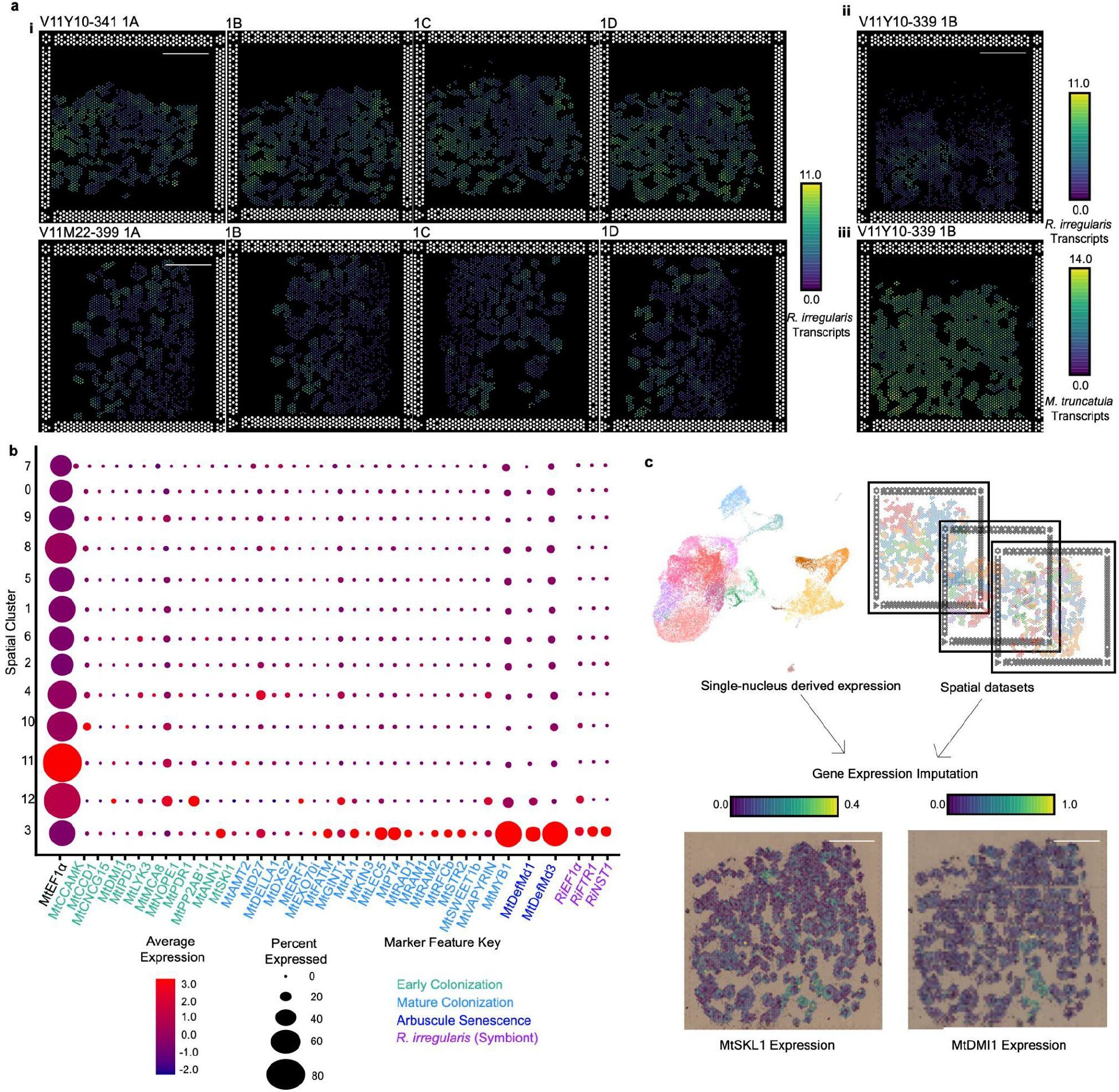
Transcriptomic profiling reveals coordinated gene expression between the symbiotic partners. a, Spatial distributions of all *R. irregulari*s and *M. truncatula* transcript expression from 9 unique capture areas containing lateral root cross-sections from *M. truncatula* plants at 28 dpi (scale bar = 1 mm, scale = log2(UMI counts)). i, 8 unique inoculated capture areas exhibiting the spatial distribution of all *R. irregularis* transcripts within the roots. ii, A single inoculated capture area showing differing spatial distributions of all *R. irregularis* transcripts (top image) and *M. truncatula* transcripts (bottom image). b, Dotplot of *M. truncatula* and *R. irregularis* housekeeping genes along with genes known to be involved in different stages of the symbiosis utilizing hierarchical clustering within the integrated mycorrhizal spatial object. c, Visualization of gene expression imputation from single-nuclei mycorrhizal integrated dataset for two lowly expressed genes, MtSKL1 and MtDMI, within a single Visium Spatial Gene Expression Capture area.

**Figure 3:**
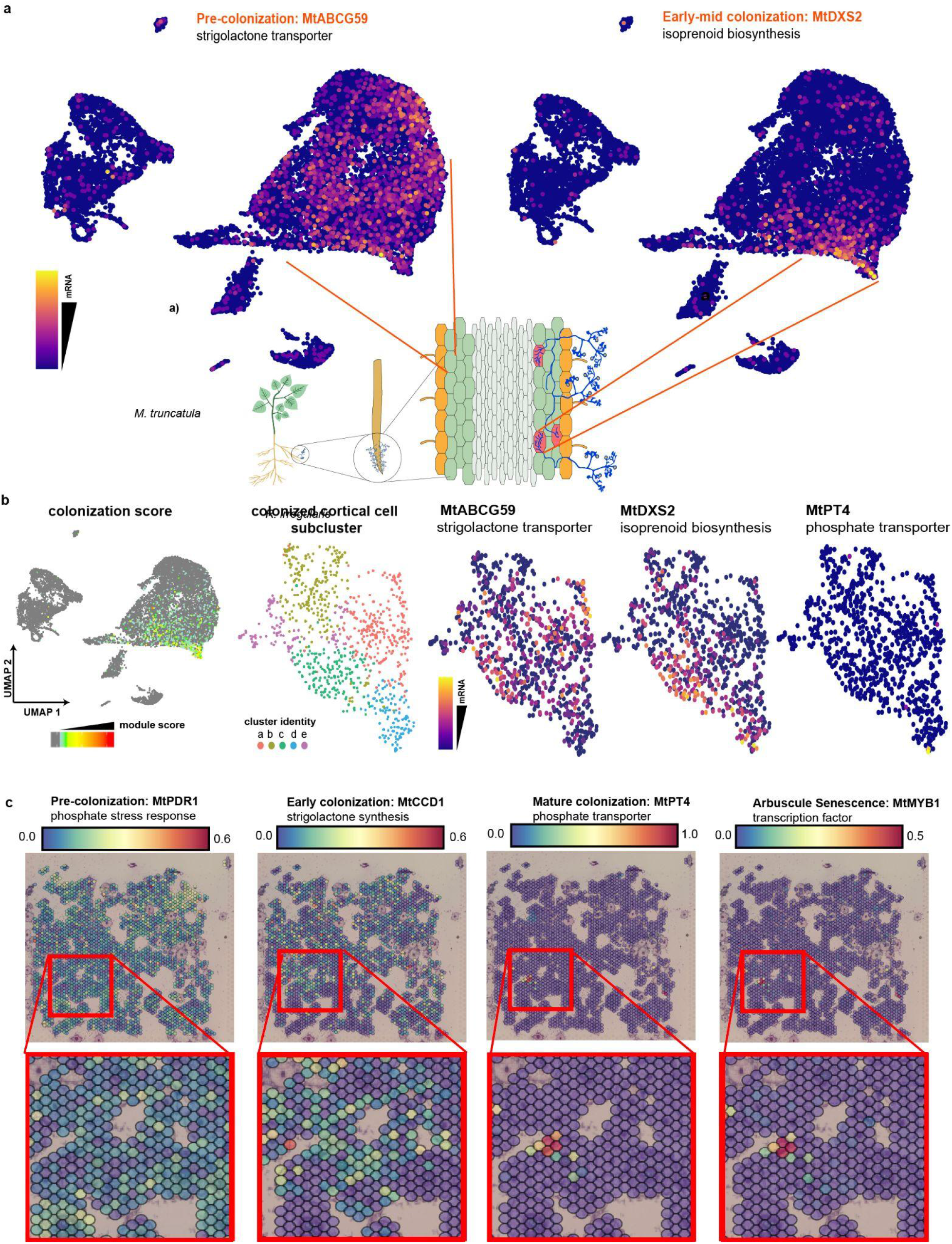
Analyzing colonization stage-specific gene expression. a, Expression of canonical pre-and post-AM marker genes shown in a UMAP plot of single nuclei dataset, colored by normalized mRNA counts. (scale = log(UMI counts + 1)). b, Module scores based on average expression level of a list of known AM marker genes; cells with module scores in the 95th percentile were selected as ‘colonized cortex cells’ and clustering was performed on this subset. Three AM marker genes known to be expressed at different colonization stages are expressed in different subclusters of the colonized cluster. c, Spatial feature plots of four AM marker genes known to be expressed at different colonization stages: MtPDR1, MtCCD1, MtPT4, and MtMYB1. Zoom-panels focus on voxels that switch expression profiles

**Figure 4:**
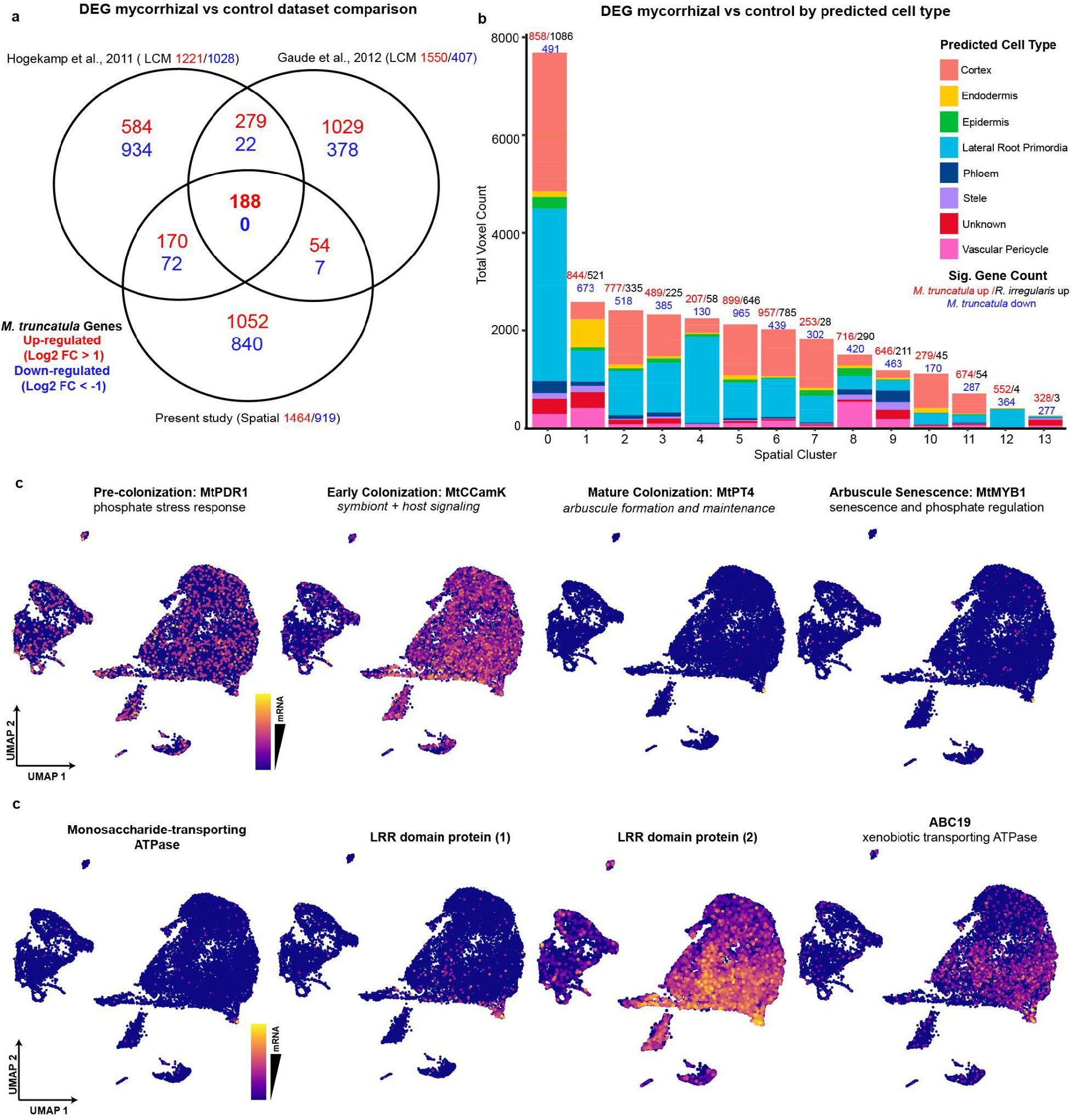
Existing and novel transcriptomic studies reveal a robust set of differentially expressed *M. truncatula* genes during the AM symbiosis. a, Venn diagram shows overlap in symbiosis-responsive *M. truncatula* genes with a log-fold change > or < 1 between the spatial dataset rom this study and the two previously published laser-capture microdissection RNA-seq studies from Guade et al. 2012^15^ and Hogekamp et al. 2013^14^. b, Breakdown of predicted cell types represented in each cluster within the integrated spatial dataset in terms of number of voxels. Differentially expressed genes between mycorrhizal and control treatments are shown above each bar for each cluster, with *M. truncatula* up regulated counts in red and downregulated counts in blue. Significant *R. irregularis* gene counts are expressed in black. c, (upper) Expression of four canonical pre-and post-AM marker genes shown in a UMAP plot of single nuclei dataset. (lower) Expression of four genes enriched in the colonized cortex cell cluster of this dataset which have yet unknown roles in AMF colonization, shown in a UMAP plot of single nuclei dataset. UMAP plots colored by normalized mRNA counts (scale = log(UMI counts + 1)) depending on the marker gene, thus reflecting the colonization stage.

### Spatial transcriptomics reveals overlap in colonization-specific transcriptomic signatures of the plant and fungus

To identify any features associated with AMF colonization within our spatial transcriptome datasets, we performed dimensionality reduction and clustering of all voxel transcriptome profiles from both AMF-and mock-inoculated spatial capture areas (Fig. S5). Due to the relatively low resolution of the Visium Spatial Gene Expression slide (each 55 µ*m* voxel could contain between 1 and 5 root cells), we refrained from assigning cell type identity to the various spatial clusters, as voxels are most likely representing heterogeneous cell groups. Instead, we focused on identifying clusters of voxels within the mycorrhizal dataset that represent sites of AM colonization. To do so, we compiled a list of AM-responsive marker *M. truncatula* and *R. irregularis* genes from the literature (Table S3) that corresponded to the various stages of AM colonization: early colonization, mature colonization, or arbuscule senescence. We analyzed the expression of these marker genes across the 13 unique spatial clusters in comparison to that of *ELONGATION FACTOR 1-ALPHA (MtEF1α)* which was reported to be stably expressed during the AM symbiosis^46^. Spatial clusters 3 and 12 showed high specific expression of the AM marker genes from both species and thus were deemed “AM-responsive” (Fig. 2b), with spatial cluster 12 showing higher expression of early marker genes and spatial cluster 3 showing higher expression of mature and senescence colonization stage markers. Of note, there were a few AM marker genes which were detected within the single-nuclei dataset but were either missing or lowly expressed within the spatial datasets. This could be attributed to the lower detection efficiency of unbiased transcriptomic methods as compared to probe-based capture^47^. In such cases, it was possible to integrate the single-nuclei and spatial datasets and impute the missing expression values using the TransferData() function within Seurat^19^. Two examples of this imputation are shown in Fig. 2c, with the expression distribution of two AM-marker genes initially missing from the dataset, *DOES NOT MAKE INFECTION 1 (MtDMI1)* and *SICKLE 1* (*MtSKl1)*, imputed onto a mycorrhizal capture area.

### Single nuclei and spatial transcriptomics reveal colonization-stage specific gene expression

Genes known to be involved in specific stages of AM colonization were selected as markers to determine the distribution of cells in different stages of arbuscule formation. Low-phosphate conditions stimulate the interaction between plant and fungus, which results in the secretion of strigolactones from the root cortex cells. The ABCG59 transporter is known to secrete strigolactones into the extracellular space, and mRNA transcripts of *MtABCG59* are enriched in cells throughout the cortex clusters 0, 1, 2, 4, 6, 8, 9, 14, and 16, which we combined into one cluster for further analysis (Fig. 3a), representing cells in a pre-colonization state. Cluster 14, which specifically expressed *1-DEOXY-D-XYLULOSE 5-PHOSPHATE SYNTHASE* (*MtDXS2)* transcripts, likely represents cells undergoing active AM symbiosis, as *MtDXS2* is required for the methyl-D-erythritol phosphate pathway-based isoprenoid production to sustain AM colonization (Fig. 3a). In order to determine if a developmental gradient exists within the AM symbiosis cluster (cluster 14), we defined an expression module of known marker genes which were expressed in this cluster and scored all cells according to their expression of this AM-colonization profile. We then selected cells in the 98th percentile of this AM module score and re-clustered them into five sub-clusters (Fig. 3b). Based on the enrichment of marker genes known to be expressed at different stages of colonization, sub-clusters a, b, and e likely represent earlier stages, as these cells are enriched for *MtABCG59* transcripts. Clusters c and d may represent later stages of colonization based on the enrichment of *MtDXS2* transcripts, and the cells at the edge of cluster d are likely at the most advanced stage of colonization, as they are the only cells in the single cell datasets that contain high levels of *MtPT4* mRNA, which is necessary for the transfer of phosphate from AM fungus to plant (Fig. 3b).

We were also able to visualize the spatial dynamics of colonization within the mycorrhizal capture areas by tracking the distribution of known AM marker genes expressed at different stages of colonization. By analyzing the expression of a pre-colonization marker, *MtABCG59* (homolog of *PaPDR1*)^48^, an early colonization marker, *MtCCD1* (*CAROTENOID CLEAVAGE DIOXYGENASE* 1)^49^, a mature colonization marker, *MtPT4*^9^, and *MtMYB1* (*MYB-LIKE TRANSCRIPTION FACTOR 1*)^31^, a gene expressed throughout arbuscule development involved in arbuscule senescence, we could identify voxels and thus overlaying individual root cross sections that were experiencing distinct stages of colonization within the capture area (Fig. 3c).

We used known marker genes to show the developmental trajectory of AM colonization across the dataset, starting with the canonical phosphate stress response gene *MtPDR1(PLEIOTROPIC DRUG RESISTANCE 1)*, which is enriched throughout the cortex cluster and represents cells in a pre-colonization state. *MtCCaMK (CALCIUM AND CA2+/CALMODULIN-DEPENDENT PROTEIN KINASE)* is involved in host-symbiont signaling in early stages, and is enriched in the cells closer to the colonized cluster. Canonical phosphate transporter *MtPT4* is enriched in a subset of cells at the furthest edge of the colonized cluster and *MtMYB1* is also enriched in these distal cells (Fig. 4c, upper panel).

Performing differential gene expression (DEG) analysis between cluster 14 and all other cortex clusters combined, we found 258 genes enriched in the AM symbiosis cluster (logFC> 0.25, adjusted p < 0.01). We found known marker genes for AM symbiosis among these upregulated genes, along with many genes which were not known to be involved in AM signaling and genes of unknown function. For example, a monosaccharide transporting ATPase (Medtr8g006790) may be used by the plant to provide sugars as nutrients or signaling molecules to the fungus, several LRR-domain proteins (Medtr6g037750, Medtr3g058840) which may be involved in host-symbiont signaling, and *MtABC19*, which a xenobiotic-transporting ATPase (Medtr3g093430) not previously known to be involved in AM symbiosis (Fig. 4c, lower panel).

### Existing and novel transcriptomic studies reveal a robust set of differentially expressed M. truncatula genes during the AM symbiosis

Differentially expressed genes (DEGs) across the 17 spatial datasets were identified by comparing the expression levels under mycorrhizal and control treatments using the DESeq2 package as described in methods. Our analysis resulted in the identification of 2,383 *M. truncatula* transcripts as AM-responsive (Fig. 4a). Among the 2,383 significantly expressed genes, 1464 were upregulated (log2 FC > 1.0) and 919 were downregulated (log2 FC< 1.0). Two previous laser-capture microdissection-based transcriptomic analyses by Guade et al. in 2012 and Hogekamp et al. in 2013 revealed similar numbers of up-regulated vs. downregulated genes in response to mycorrhizal treatment, with 188 genes significantly upregulated across all three datasets which we refer to as “robust” AM-responsive genes (Fig. 4a). No genes were found to be significantly downregulated in all three datasets, although there were common downregulated genes across pairs of datasets. This set of robust AM-responsive genes includes many characterized AM-marker genes critical to the symbiosis, such as *MtMYB1*, *MtPT4*, and *MtRAD1*^7, 30, 50^. However, many of the 188 genes are uncharacterized and remain to be investigated for their role in the AM symbiosis.

As mentioned previously, the 55 μm resolution of the Visium Spatial Gene Expression platform does not allow for single-cell resolution. However, we utilized our annotated sNuc-RNAseq datasets and spatial datasets to predict the proportions of each cell type represented in each spatial cluster (Fig. 4b) as voxels are likely to contain various cell types simultaneously. The most well represented cell-type predicted was cortex cells, which aligns with our sNuc-RNAseq data. Cortex cells also make up the majority of cell area in 2-dimensional root cross-sections. We were also able to see differences in the proportion of certain cell types between spatial clusters, such as the high proportion of endodermis represented in spatial cluster 2, and the high proportion of vascular pericycle represented in spatial cluster 8. Notably, we saw a relatively high proportion of voxels identifying as lateral root primordia, which may be attributed to the high amount of mRNA that is captured from meristematic root tissue on the capture area. We also observed large differences in the amount of DEGs between the individual spatial clusters, especially in the number of significant *R. irregularis* genes. For example, spatial cluster 0, which had the highest proportion of voxels predicted as cortex cells, also exhibited the highest number of significant *R. irregularis* genes, while spatial clusters 12 and 13, which were most represented by lateral root primordia and unknown cell identities. This may indicate that even at lower resolution, the Visium technology is able to discern between groups of root cells at varying levels of AM symbiosis involvement.

### Functional enrichment analysis depicts a marked symbiotic response within M. truncatula

To investigate our dataset and the robust set further, we performed gene and protein functional classification for both using the PANTHER Classification system (www.pantherdb.org)^51^ (File S7). Functional enrichment analysis of parent gene ontology categories was performed for biological processes (Fig. S6a), molecular functions (Fig. S6b), and cellular components (Fig. S6c) represented within the list of significantly upregulated *M. truncatula* genes (panels i) and the robust set (panels ii). As expected, for the “arbuscular mycorrhizal association” biological process, we saw a >20 fold enrichment within our spatial dataset and a >80 fold enrichment within the robust set. We also observed a high enrichment (>100 fold) of the “response to symbiotic bacterium” within the robust set, which may be indicative of genes common to both the AM and rhizobial symbioses. Interestingly, the biological process “proline catabolic process to glutamate” and the molecular function “proline dehydrogenase” showed a >20 fold enrichment (Fig. S5a and b). Several studies have observed altered proline levels specifically under drought stress in AM-treated plants^52, 53^, which is thought to confer some protection against the stress, although the exact mechanism remains unclear. As these plants were not experiencing significant drought stress, it may indicate a different stress response or protective mechanism at work in the root cells.

AM fungi are known to convert soil inorganic phosphate (Pi) into inorganic polyphosphate (polyP) and are thought to be able to rapidly accumulate and translocate polyP within their hyphae^54^. AM fungi also have the ability to depolymerize polyP via fungal endopolyphosphatases and release phosphate from the arbuscule in order to transfer this phosphorus into host plant cells across the periarbuscular membrane^55^, although the mechanism for this export remains unclear. There is plenty of evidence to support the hypothesis that the majority of this export occurs via the transport of Pi across the apoplastic space and subsequent uptake by the plant via Pi transporters. However, there is a growing amount of evidence that suggests polyP may be directly exported to the apoplastic space and then hydrolyzed by the plant itself^58^. A study conducted by Nguyen and Saito in 2021 provided evidence that fungal-derived polyP and plant-derived phosphatases had opposite localizations in mature arbuscules, indicating that plant phosphatase activity could be responsible for fungal-derived polyP^56^. Surprisingly, “exopolyphosphatase activity” exhibited the highest enrichment of all molecular functions, at >30 fold in our dataset. This adds support to the hypothesis that polyphosphatase activity by the plant plays a larger role in phosphorus export than previously anticipated. Further analysis of the transcripts that were characterized by this molecular function is encouraged.

Lastly, the cellular components enriched in our spatial dataset and the robust dataset were as expected composed of cell components involved in the AM symbiosis, such as the periarbuscular membrane and casparian strip. Of note, the robust dataset showed a >100 fold enrichment for the “periarbuscular membrane” category and was able to capture 100% of the *M. truncatula* genes assigned that functional category within the genome (Fig. S6c). Overall, the gene ontology annotation and functional enrichment analysis confirmed a strong symbiotic signature within our mycorrhizal dataset and revealed interesting functional categories for further investigation.

### Spatially-resolved R. irregularis transcripts reveal novel AM-specific gene expression patterns

Numerous whole-root and single-cell RNA-seq studies have been conducted on *M. truncatula* in symbiosis with *R. irregularis* utilizing a broad spectrum of cell isolation and transcriptomic techniques. However, the majority of data collected has come from the plant species *M. truncatula* while the fungal transcriptome remains elusive and resistant to many mRNA capture technologies. One major effort by Tisserant et al. in 2011 provided a glimpse into the fungal symbiont transcriptome for the first time by sequencing RNA from germinated fungal spores, extra and intraradical mycelium, and arbuscules via Sanger and 454 sequencing as well as microarray profiling^57^. In addition, two laser-capture microdissection-based transcriptomic analyses conducted by Guade et al. in 2012 and Hogekamp et al. in 2013 were able to capture numerous fungal transcripts hybridized to the Affymetrix Medicago GeneChip array within mycorrhizal roots^13, 39^ and further improved our understanding of the symbiotic response from *R. irregularis*. Our spatial transcriptomics approach built upon this existing research by providing the first spatially-resolved dataset of simultaneously captured plant and fungal transcriptomes during AM symbiosis. We detected the expression of 12,104 unique fungal transcripts across the nine mycorrhizal capture areas (Supp. File 3) and identified 14 fungal transcripts conserved between our study and the two laser-capture microdissection based analyses (Fig. 5a). Of the 12,104 expressed fungal genes, differential gene analysis revealed 4,817 to be significantly upregulated in mycorrhizal capture areas as compared to datasets from non-inoculated samples (p-value=0.05, log2 FC >1.0). We were also able to correlate fungal expression distributions across the capture area with the existence of arbuscules in the tissue, with root cross-sections that displayed high numbers of arbuscules in the cortex also displaying high expression of *R. irregularis* transcripts in the underlying voxels (Fig. 5b). These spatial foci of *R. irregularis* were also transcriptomically distinct, as *RiEF1*α *(ELONGATION FACTOR 1-ALPHA)*, a housekeeping elongation factor gene^58^, *RiFTR1*, an iron ion permease^59^, and a highly-expressed but uncharacterized feature, RIR_103690, all show high and specific expression to the AM-responsive cluster 3 in UMAP space, indicating that the *R. irregularis* transcript expression in those voxels is symbiosis-specific (Fig. 5c). Lastly, despite large improvements in the *R. irregularis* genome assembly, gene annotation and characterization is still limited. We conducted gene ontology analysis with the Blast2Go software^60^ (Blast2Go, https://www.biobam.com/blast2go)^55^ for the top 100 expressed *R. irregularis* transcripts, and though many transcripts could not be annotated, a large number of terms associated with membrane localization, as well as protein and lipid metabolism were enriched (Fig. S5d). This dataset of more than 12,000 symbiosis-specific spatially-resolved *R. irregularis* genes and corresponding images of fungal structures within the root cross sections is the first of its kind and holds immense potential for in-depth characterization of symbiosis-responsive fungal genes.

**Figure 5:**
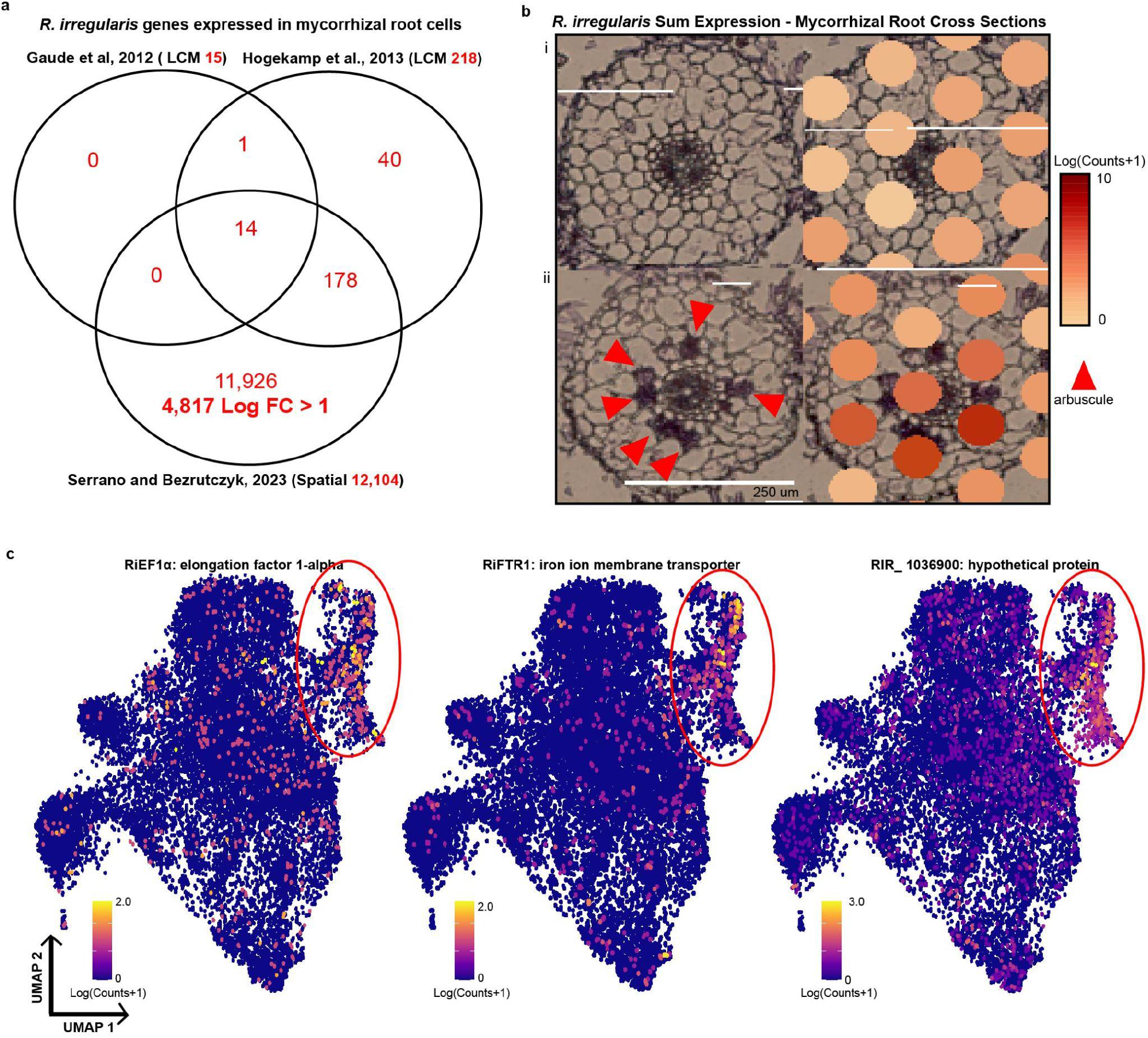
Spatially-resolved *R. irregularis* transcripts reveal novel AM-specific gene expression patterns. a, Venn diagram shows overlap in total expressed *R. irregularis* genes between the spatial mycorrhizal dataset from this study and the two previously published laser-capture microdissection RNA-seq studies from Guade et al. 2012^15^ and Hogekamp et al. 2013^14^. b, Brightfield images of a single root cross-section within the spatial gene expression capture area (left) and overlapping spatial distributions of all *R. irregularis* transcript expression (right) visualized in Loupe Browser. i, Root cross-section from a mycorrhizal capture area that is lacking recognizable fungal structures shows low overall expression of *R. irregularis* transcripts. ii, Root cross-section from a mycorrhizal capture area that contains visible arbuscules (red arrows) shows high overall expression of *R. irregularis* transcripts, particularly around arbuscules (scale bar = 250 um, scale = log(UMI counts + 1)). c, Representative feature plots of a housekeeping gene, RiEF1a, a known AM-marker gene, RiFTR1, and a previously undescribed gene, RIR_103690, that has high expression across AM-associated cluster in the integrated mycorrhizal spatial dataset (scale = log(UMI counts + 1)).

## Discussion

Advances in single-cell transcriptomic technology have transformed the field of molecular genetics by allowing for cell-type specific transcriptomic analysis and, more recently, the conservation of the tissue context, adding in another dimension to the data^61^. Here, we combined two complementary technologies, single-nucleus and spatial RNA-seq, to construct the first spatially-resolved single-cell resolution integrated map of a multi-kingdom symbiotic interaction to date. We successfully adapted the spatial transcriptomics platform for use with plant roots and utilized unbiased, transcriptome-wide mRNA capture to detect mRNA from two species simultaneously. In addition, we effectively identified cell type-specific responses to the AM symbiosis using single-nuclei RNA-seq and were able to integrate data from both approaches to discover novel AM-responsive *M. truncatula* and *R. irregularis* transcripts.

Differential gene expression analyses between the AM-colonized cortex cell cluster and other cortex cells of the inoculated datasets revealed 258 genes which are significantly upregulated in the colonized cortex cells. These genes were enriched for GO terms related to fungal symbiosis, terpene synthesis pathways, and transport proteins. We found 17 different cytochrome P450-like proteins, whose roles in plant-microbe interactions include hydroxylation of fatty acids to provide fungal symbionts with carbon, as well as synthesis of terpenes for plant defense^62^. There were 19 leucine-rich repeat (LRR) domain-containing proteins, which are critical for signaling cascades resulting from microbe-associated molecular pattern recognition, and while typically associated with plant-pathogen interactions, are also upregulated in plant-AM fungal symbiosis^63, 64^. ABCG transporters such as *MtABCG59* are known to secrete strigolactones in response to phosphate stress^48^, and were found enriched in the colonized cortex cluster (CCC). Additionally, several xenobiotic, sugar, and amino acid transporters were also significantly upregulated in the colonized cortex cluster, many of which may play unknown roles in AM symbiosis. The identification of known and previously unknown transcripts within the colonized cortex cell cluster presents a community resource for the characterization of novel AM-associated genes.

Single-nuclei RNA-seq provides increased resolution and throughput, for the identification of cell-state and cell-type responses to external treatments^65^. Single-nuclei RNA-seq eliminates the hurdle of protoplasting cells and instead relies on a quick and simple nuclei extraction protocol which is adaptable to diverse tissue types and species. Ideally, this technique would have allowed us to obtain cells at various stages of arbuscule development– however, on disadvantage of this protocol relative to LCM is that we were not able to visually confirm cortex cells with arbuscules, resulting in a low proportion of arbusculated cortex cells in our sNucRNAseq datasets. snRNA-seq datasets largely reconstruct the cell types covered in single cell datasets, although with lower gene diversity and transcript counts per cell, but have the advantage of reducing bias in sampling due to a more uniform profiling of all cell types, including those more difficult to enzymatically release from their cell walls. For example, cells in which arbuscules form have highly ramified cell membranes, which may be difficult to recover after enzymatic digestion due to their increased surface area, further limiting our access to an already limited group of cells within the larger root cell population. The process of isolating nuclei from root tissue is also faster and results in the upregulation of fewer stress-related genes^66^. Another potential advantage of using nuclei over protoplasts was the possibility of obtaining fungal nuclei in the same assay. *R. irregularis* consists of multinucleate hyphae in a large syncytium, and would therefore not be amenable to protoplast formation. However, even in batches of nuclei which were not subjected to gated flow-sorting, we were unable to recover a significant number of fungal transcripts and obtained no defined fungal nuclei, possibly because fungal nuclei were not captured or were destroyed by our plant nuclei extraction protocol. We were able to overcome this limitation of snRNA-seq by supplementing with results from the spatial transcriptomics assay.

Spatial transcriptomics allowed for the preservation of the tissue context during sequencing and for a side-by-side comparison of gene expression and cellular features. However, limitations of this technology posed several challenges. The main limitation of the majority of transcriptome-wide unbiased spatial transcriptomic technologies is low spatial resolution (large voxel size and inter-voxel distances). Higher resolution is possible in probe-based capture spatial technologies, although researchers are limited to analyzing a set of known genes for which probes have been specifically designed^58^. As the resolution from our study is effectively 50µm, cell-type identity during clustering analysis cannot be accurately defined due to each voxel containing 1-5 root cells (with adjacent root cells representing very different cell types). Increasing resolution of these technologies leads to lower capture efficiency due to the lower number of reads that can be captured by fewer primers^67^. There is clearly a need for a high-resolution spatial transcriptomic technology that allows for unbiased mRNA capture from intact tissue.

Our study identified thousands of *R. irregularis* genes expressed in mycorrhizal *M. truncatula* root cells and provided evidence that this gene expression was specific to cells that were AM-responsive based on *M. truncatula* AM-marker gene expression. *R. irregularis* is a widespread symbiont of many diverse plant species and is widely used in commercial agriculture due to its ability to uptake P and N from soils^68^. It is a model species for the AM symbiosis and fungal biology, making it one of the most studied fungi^69^. However, despite the agricultural and biological importance of this fungus, functional annotation and characterization of its genes has been a slow process due to the difficulty in genetic manipulation of the organism^70^. Recent advances in CRISPR-Cas9^71^ gene editing for fungal species and continued efforts in creating pure cultures of AM fungi^72^ will hopefully lead into a new chapter of AM symbiotic research in which the fungal partner is the focus. The many previously uncharacterized genes differentially expressed between mycorrhizal and control treatments within our study, particularly within the 188 “robust” gene list common to three distinct RNA-seq datasets, present as excellent targets for CRISPR-Cas9 modification. Expression modulation of these genes within a symbiotic context could reveal novel aspects of plant-AM fungal interactions.

Moving forward into the new era of transcriptomics in plants, we strongly believe that the scientific community at large would benefit from increased efforts in protocol and resource sharing. Because these approaches yield large, complex datasets that will prove to be useful to many diverse research groups and applications, we anticipate that efforts towards building community resources will be crucial to the advancement of transcriptomic research in plants. Resources such as the Plant Cell Atlas^73^, an initiative aimed at setting standards for creating and integrating plant single-cell data sets, are a good starting point for the community to develop a framework for the standardization of data reporting and sharing.

## Methods

### Plant growth and inoculation

Seeds of *Medicago truncatula* Gaertn, cv Jemalong A17 (Noble Foundation) were scarified in concentrated sulfuric acid for 5-10 min and rinsed with distilled water, sterilized in 3.75% sodium hypochlorite solution, then rinsed five times with sterile distilled water and placed on 1⁄2 Murashige and Skoog medium, 1% agar plates at 4C or RT for 48h. Sand cones were prepared as follows: 8.25 inch cone-tainers (Stuewe and Sons) with 1 cm^3^ rock wool at base, filled up to 12.7 cm with autoclaved calcined clay (Turface Athletics MVP 50), followed by 2.5 cm of autoclaved horticultural sand (American Soil and Stone) and topped with 2.5 cm fine play sand (SAKRETE). To inoculate seedlings with *Rhizophagus irregularis* (Błaszk., Wubet, Renker and Buscot) C. Walker and A. Schüßler: 50 mL Agtiv Field Crops liquid mycorrhizal inoculant (PremierTech, Rivière-du-Loup, Québec, Canada) spores were captured on a 40 µm filter, rinsed with distilled water and resuspended in 50 mL distilled water. 1 mL of resuspended spores was applied to the horticultural sand layer and an additional 300 µL applied to the fine play sand layer. Germinated seedlings were transplanted to the top fine sand layer of inoculated and non-inoculated sand cones. Plants were grown in 22-24 °C 16 h day/ 8 h night, with 300 μmol m^-2^ s^-1^ light intensity, and 60% relative humidity. Plants were watered daily and fertilized twice a week with 1/2X Hoagland’s medium modified with 20 μM phosphate to stimulate AM colonization.

### Colonization assessment

AM colonization in WT roots at 21, 28, or 38 dpi was visualized via staining of AM chitin using 2.5 μg/mL wheat germ agglutinin (WGA) Alexa Fluor 488 (Thermo Fisher Scientific) in 1X Phosphate-Buffered Saline solution (pH 7.0). Briefly, roots collected from the fine sand layer were rinsed, fixed in 50% ethanol for 30 minutes, and cleared in 10% KOH at 65 °C for 48 h. Cleared roots were neutralized with 0.1 M HCl and stained with WGA-488 in 1X PBS at 4 °C for 24 h prior to imaging. Colonization was quantified using the Trouvelot method^74^ on a Leica DM6B fluorescence microscope using five biological replicates for each treatment (Fig S2a).

### Quantitative real-time PCR of target genes

To quantify expression of target genes, 100 mg of roots from the fine sand layer were flash frozen in liquid nitrogen. Total RNA was extracted using the RNeasy Plant Mini Kit (Qiagen) and corresponding DNase. cDNA synthesis was conducted using the SuperScript™ IV Reverse Transcriptase (Thermo Fisher Scientific) from 500 ng of total RNA and qPCR was conducted from cDNA diluted 1:5 using the PowerUp SYBR Green Master Mix (Thermo Fisher Scientific). A 200 nM primer concentration and the following protocol was used for qPCR for all targets: 2 min at 50 °C and 2 min at 95°C, followed by 39 repeats of 15 s at 95 °C, 15 s at 60 °C and 1 min at 72 °C, and ending with 5 s at 95°C. A melting curve (55–95 °C; at increments of 0.5 °C) was generated to verify the specificity of primer amplification. Five biological replicates and three technical replicates of all targets (*MtPT4*, *RiTUB*, and *MtGINT1*) were quantified for gene expression levels relative to the housekeeping gene *MtEF1α* using the ΔΔCT method (Fig. S2b). All primer sequences used for qPCR can be found in Table S1.

### Nuclei and bulk root tissue RNA profiling

*M. truncatula* roots from three plants from each treatment were harvested at 21, 28, and 38 days post inoculation; 150 mg of roots grown in the inoculated fine sand layer were either flash frozen in liquid nitrogen, or nuclei extraction was performed up to the 20 µm filtration step, and then flash frozen. RNA was extracted using the RNeasy Plant Mini Kit (Qiagen). Library preparation and sequencing were performed at the QB3 UC Berkeley Genomics Core Sequencing Facility.

### Nuclei extraction and sequencing

*M. truncatula* roots from three plants per condition were harvested at 21-, 28-, or 38-dpi. 150 mg of roots growing in inoculated fine sand layer was weighed out and placed in the lid of a petri dish and chopped rapidly with a razor blade for three minutes in 600µL NIBAM: 1x NIB (Sigma CELLYTPN1-1KT), 4% BSA, 1 mM DTT, 0.4 U/µL Superase RNAse inhibitor (Sigma), 1:100 Protease Inhibitor Cocktail for plant tissues (Sigma). NIBAM-root slurry was strained through 40 µm and 20 µm filters (CellTrics). SYBR Green (1:10000) was used to visualize nuclei during purification on the Influx Flow Cytometer. 20,000 nuclei were sorted into 19 µL of ‘landing buffer’ (phosphate buffered saline with 0.4U/µL Superase RNAse inhibitor) with a final volume of 43 µL. DAPI was applied to 2 µL of nuclei suspension to evaluate the quality of nuclei on a Leica AxioObserver at 20x magnification. The remaining 41 µL was mixed with 10x Genomics Chromium RT Master Mix with no additional water added and loaded onto a Chromium Chip G, and thereafter the standard manufacturer’s protocol was followed (V3.1 Dual Index). Twelve cycles were used for cDNA amplification, and the completed cDNA library was quantified using an Agilent Bioanalyzer. Sequencing was performed at the QB3 UC Berkeley Genomics Core Sequencing Facility on a single NovaSeq SP lane with the sequencing parameters: 28 bp (read 1 length), 10 bp (index 1 length), 10 bp (index 2 length), 90 bp (read 2 length), or at Novogene (Sacramento, CA) using the sequencing parameters: 150bp (read 1 length), 10 bp (index 1), 10 bp (index2) and 150 bp (read 2 length).

### Tissue preparation for spatial transcriptomics

Spatial transcriptomics was performed with the Visium Spatial Gene Expression platform from 10X Genomics. Harvested plant roots were rinsed with deionized H_2_O and cryopreserved in OCT (Optimal Cutting Temperature compound) via submerging of OCT-embedded molds into a dewar of isopentane chilled liquid nitrogen for even freezing. Cryomolds of roots were stored at −80 °C until cryosectioning. Cryosectioning was performed on an Epredia™ CryoStar™ NX70 Cryostat with a blade temperature of −14 °C and a sample head temperature of –12 °C with a section thickness of 16 µ*m*. Cryosections were placed onto the surface of the chilled Visium Spatial Gene Expression slide and adhered to the slide using heat from the sectioner’s finger placed on the back surface of the capture area. Prepared slides were stored at −80 °C prior to processing for 10x Visium spatial transcriptome sequencing according to the manufacturer’s instructions with the following modifications: First, cryosections were stained using an incubation of 0.05% Toluidine Blue-O in 1X PBS for 1 min and rinsed 3X with 1X PBS. Second, A pre-permeabilization step was added as suggested by Giacomello et al.^18^ The pre-permeabilization mix for each slide (48 µl Exonuclease I, 4.5 µl of Bovine Serum Albumin, and 428 µl of 2% PVP 40) was then prepared, and 70 µl was pipetted into each well. Pre-permeabilization occurred for 30 min at 37 °C after which the manufacturer’s protocol for tissue permeabilization was followed. Tissue permeabilization occurred for 12 min based on the results of the manufacturer’s Tissue Optimization protocol (Fig. S1).

### Data processing

Cellranger and Spaceranger software (10X Genomics, Pleasanton, CA) were used to preprocess single nuclei and spatial transcriptomic sequencing libraries respectively. A formatted reference genome was generated using Cellranger or Spaceranger’s “mkref” function using the *Medicago truncatula* MedtrA17_4.0^75^ whole genome sequence and annotation and the *Rhizophagus irregularis* Rir_HGAP_ii_V2 (DAOM 181602, DAOM 197198)^76^ whole genome sequence and annotation using default parameters. Single-nuclei and spatial reads were aligned to the genome references using the “count” function in Cellranger and Spaceranger software packages (10x Genomics, Pleasanton, CA), respectively, which use STAR^77^ as the alignment engine Brightfield tissue images were aligned to the spatial capture area fiducial frame and voxels corresponding to overlaying tissue were manually selected for all capture areas in Loupe Browser (10X Genomics).

### Data analysis

Data analysis for both the single nuclei and spatial data was performed using the Seurat R package^19, 43–45^. Raw feature and UMI counts for all datasets are displayed in File S1. All scripts used for data analysis are available on GitHub at https://github.com/kserrano109/Medicago_Rhizophagus_RNAseq. Data availability on NCBI GEO (Gene Expression Omnibus) is available at https://www.ncbi.nlm.nih.gov/geo/query/acc.cgi?acc=GSE240107.

### Filtering and normalization

For both single nuclei and spatial datasets, normalization and scaling were performed using the SCTransform R function in Seurat prior to clustering. Metrics used for filtering of the data during quality control steps can be found in File S1.

### Principal components analysis and K-means clustering

Principal components analysis was performed on both snRNA-seq and spatial RNA-seq datasets using the RunPCA() function in Seurat with the “SCT” assay specified. The FindNeighbors() function was applied to construct a Shared Nearest Neighbor (SNN) graph for the data using the first 30 principal components. Clustering was performed using the FindClusters() function which utilizes the SNN graph from the previous step. Finally, the RunUMAP() function was utilized to construct the Uniform Manifold Approximation and Projection dimensionality reduction and visualize the dataset in two dimensions.

### Integration of replicate datasets

Replicate capture areas from each treatment (mycorrhizal or mock-inoculated) were integrated into a single Seurat spatial object using the IntegrateData() function. First, since we utilized the SCTransform method for normalization, we applied the PrepSCTIntegration() function with all features specified as the features to integrate. We then identified a set of integration anchors with the FindIntegrationAnchors() function. Finally, within IntegrateData(), we specified SCT as the normalization method and set all features specified as the features to integrate. Principal component analysis and dimensionality reduction was performed on the integrated objects in the same manner as the individual objects with the following adjustments: 1) number of PCs = 30, 2) metric = cosine, and 3) resolution was set to 0.5 for clustering. UMAP plots for all datasets were created using the DimPlot() function and are displayed in Fig. S4 (single-nuclei) and Fig. S5 (spatial).

### Differential gene expression

Differentially expressed genes (upregulated and downregulated) between the mycorrhizal and control integrated datasets for all clusters were identified using the Likelihood Ratio Test from the DESeq2^78^ package with an adjusted p-value of <0.05 and a log2 fold-change threshold of -1.0 or 1.0.

### Module score analysis

To determine which cells represented colonized cortex cells, we generated a list of genes (Table S3) which are known to be involved in colonization, and used these as input to assign a module score to each cell using the Seurat AddModuleScore function. Cells with a score above the threshold of 95th percentile were selected as ‘colonized’ cells, and subsetted to a new object for further sub-clustering analysis.

### Gene expression imputation

Using the annotated single-nuclei integrated object as a reference and the spatial integrated object as a query, we performed a data transfer using the UMI counts from the RNA assay within the single nuclei object as the reference data and stored the new data under a new assay called “imputation”. We then were able to predict gene expression within the spatial dataset using the expression values from the single-nuclei dataset by specifying the assay to “imputation” during the analysis.

### Voxel cell type proportion prediction in spatial RNA-seq

Using the annotated single-nuclei integrated object as a reference and the spatial integrated object as a query, we performed a label transfer using the cell type annotations as the reference data. The query dataset is then projected onto the PCA of the reference dataset and the labels are predicted.

### Comparison to previous datasets

We compared our dataset to two prior studies (Guade et al. 2012 and Hogekamp 2013) that significantly improved our understanding of gene expression changes during the mycorrhizal symbiosis between *M. truncatula* and *R. irregularis*. We wanted to include these datasets in our analysis to identify a core set of DEGs between the three different RNA-seq techniques and be able to compare and contrast the various methods. One major hurdle to this comparison resulted from the use of the Affymetrix Medicago GeneChip array by these two studies leading to a difference in feature IDs. We constructed an ID converter to convert between the Affymetrix GeneChip, MedtrA17_4.0, and the *M. truncatula* A17 r5.0 gene IDs for a certain locus in bulk fashion (File S5). Genes identified as common between datasets can be found in File S6.

### Gene Ontology and Functional Enrichment analysis

For *M. truncatula,* we conducted gene ontology and functional enrichment analyses utilizing the PANTHER Classification system (www.pantherdb.org)^51^. For *R. irregularis,* we conducted gene ontology analysis with the Blast2Go software^60^ (Blast2Go, https://www.biobam.com/blast2go)/.

## Supporting information

Supplementary Sheet 1

Supplementary Sheet 2

Supplementary Sheet 3

Supplementary Sheet 4

Supplementary Sheet 5

Supplementary Sheet 6

Supplementary Sheet 7

## Acknowledgements

We would like to acknowledge Christopher Gee, Lorenzo Washington, Victoria Vera, Jutta Dalton, Trevor Tivey, Bruno Guillotin, and Rihanna Hunter for their advice, expertise, and technical help. We also would like to thank the QB3 UC Berkeley Genomics Core Sequencing Facility (*QB3 Genomics, UC Berkeley, Berkeley, CA, RRID:SCR_022170)* for their help in cryosectioning and sequencing. This study was supported by the DOE Joint BioEnergy Institute (http://www.jbei.org) and the DOE Joint Genome Institute (https://ror.org/04xm1d337) supported by the U. S. Department of Energy, Office of Science, Office of Biological and Environmental Research, through contract DE-AC02-05CH11231 between Lawrence Berkeley National Laboratory and the U.S. Department of Energy. Part of the work was supported by the Laboratory Directed Research and Development program at Lawrence Berkeley National Laboratory.

## Contributions

K.S, M.B, H.S. and B.C. planned experiments; K.S., M.B., T.D., and D.G performed experiments; R.O., R.M., and A.V. provided consultation; K.S., M.B., and B.C. analyzed the data; K.S., M.B., H.S., and B.C. wrote the manuscript.

