## Supplementary Data for "Spatial Co-transcriptomics Reveals Discrete Stages of the Arbuscular Mycorrhizal Symbiosis"

**Supplementary Table 1. Primer sets used for qRT-PCR in this study.**

| **Sequence 5' → 3'** | **Primer Name** | **Gene ID** |
|---|---|---|
| TGA CAG GCG ATC TGG TAA GG | qMtEF1a - Forward | Medtr5g037343 |
| TCA GCG AAG GTC TCA ACC AC | qMtEF1a - Reverse | Medtr5g037343 |
| GGA TTC TTT TGC ACG TTC TTG G | qMtPT4 - Forward | Medtr1g028600 |
| CCT GTC ATT TGG TGT TGC AGT G | qMTPT4-Reverse | Medtr1g028600 |
| ATA CCG TGG CGA TGT CGT TC | qRiTUB - Forward | R. irregularis DAOM 197198 v2.0 - 67395 |
| GAC AAC GGT TGG GGG TTG ATA G | qRiTUB - Reverse | R. irregularis DAOM 197198 v2.0 - 67395 |

**Supplementary Table 2. Marker genes for IDing cell types for sNucRNAseq dataset.**

| **Gene ID** | **Gene Name** | **Transcript ID** | **Cell Type** | **Reference** |
|---|---|---|---|---|
| MtPHO1.1 | PHOSPHATE TRANSPORTER 1.1 | Medtr1g041695 | central cylinder/pericycle | 10.1093/plphys/kiaa016 |
| MtPHO1.2 | PHOSPHATE TRANSPORTER 1.2 | Medtr1g075640 | central cylinder/pericycle | 10.1093/plphys/kiaa016 |
| MtPHO1.3 | PHOSPHATE TRANSPORTER 1.3 | Medtr8g069955 | central cylinder/pericycle | 10.1093/plphys/kiaa016 |
| MtNOPE1 | *Major Faciltator Superfamily TRANSPORTER* | Medtr3g093270 | colonized cortex | 10.1038/nplants.2017.73 |
| MtAMN2 | *ABCB FOR MYCORRHIZATION AND NODULATION 2* | Medtr4g081190 | colonized cortex | 10.1094/MPMI-02-21-0036-R |
| MtAMN3 | *ABCB FOR MYCORRHIZATION AND NODULATION 3* | Medtr8g022270 | colonized cortex | 10.1094/MPMI-02-21-0036-R |
| MtDXS2 | *1-deoxy-D-xylulose 5-phosphate synthase* | Medtr8g068265 | colonized cortex | 10.1016/j.cub.2017.03.003 |
| MtRAM1 | *REDUCED ARBUSCULAR MYCORRHIZA1* | Medtr7g027190 | colonized cortex | 10.1016/j.cub.2017.03.003 |
| MtVPY | *VAPYRIN* | Medtr1g089180 | colonized cortex | 10.1111/j.1365-313X.2010.04415.x |
| MtSUNN | SUPER NUMERIC NODULES | Medtr4g070970 | companion cells | 10.1093/jxb/eraa193 |

| | | | | |
|---|---|---|---|---|
| AtSUC2 homolog | *SUCROSE PROTON SYMPORTER2* | Medtr1g096910 | companion cells | 10.1093/plcell/koaa060 |
| AtFTIP1 | *FT INTERACTING PROTEIN1* | Medtr0291s0010 | companion cells | 10.1093/plcell/koaa060 |
| MtIFS1 | *ISOFLAVONE SYNTHASE1* | Medtr4g088195 | cortex | 10.1093/jxb/erx059 |
| MtIFS3 | *ISOFLAVONE SYNTHASE3* | Medtr4g088160 | cortex | 10.1093/jxb/erx059 |
| MtABCG59 | *ATP-BINDING CASSETTE TRANSPORTER59* | Medtr3g107870 | cortex | 10.3389/fpls.2020.00018 |
| MtSCR | *SCARECROW* | Medtr7g074650 | endodermal cells | 10.1038/s41586-020-3016-z |
| MtPLT1 | *PLETHORA 1* | Medtr2g098180 | LRP | 10.1242/dev.120774 |
| MtPLT2 | *PLETHORA 2* | Medtr4g065370 | LRP | 10.1242/dev.120774 |
| MtPLT3 | *PLETHORA 3* | Medtr5g031880 | LRP | 10.1242/dev.120774 |
| MtPLT4 | *PLETHORA 4* | Medtr7g080460 | LRP | 10.1242/dev.120774 |
| MtFe | *FE/ALTERED PHLOEM DEVELOPMENT* | Medtr6g444980 | phloem | 10.1038/s41586-018-0839-y |
| AtPEAR1 homolog | PHLOEM EARLY DOF1 | Medtr3g077750 | phloem | 10.1038/s41586-018-0839-y |
| AtPEAR1 homolog | PHLOEM EARLY DOF1 | Medtr4g461080 | phloem | 10.1038/s41586-018-0839-y |
| MtRboHF | RESPIRATORY BURST OXIDASE HOMOLOGS | Medtr7g060540 | root hairs | 10.1021/acs.chemrestox.9b00028 |
| MtMYB07 | MYB DOMAIN PROTEIN | Medtr5g014990 | vascular | 10.1016/j.molp.2022.10.021 |
| MtMYB113 | MYB DOMAIN PROTEIN | Medtr2g096380 | vascular | 10.1016/j.molp.2022.10.021 |
| MtMYB112 | MYB DOMAIN PROTEIN | Medtr4g063100 | vascular | 10.1016/j.molp.2022.10.021 |
| MtSTP13 | *SUGAR TRANSPORT PROTEIN13* | Medtr1g104780 | vascular | 10.1111/j.1365-313X.2011.04810.x |
| MtPrx13 | PEROXIDASE13 | Medtr1g101830 | xylem | 10.1016/j.molp.2022.10.021 |
| MtLBD18 | LOB-DOMAIN PROTEIN | Medtr8g036085 | xylem | 10.1016/j.molp.2022.10.021 |

**Supplementary Table 3. List of AM Marker Genes used as an AM-specific expression module.**

| Gene Name | Gene ID | Colonization Stage | Reference |
|---|---|---|---|
| MtCCAMK | Medtr8g043970 | early | 10.1016/j.pbi.2012.04.002 |
| MtCCD1 | Medtr3g109610 | early | 10.1104/pp.108.125062 |
| MtCNCG15 | Medtr1g064240 | early | 10.1126/science.aae0109 |
| MtDMI1 | Medtr7g117580 | early | 10.1104/pp.106.086959 |
| MtIPD3 | Medtr5g026850 | early | 10.1094/MPMI-20-8-0912 |
| MtLYK3 | Medtr5g086130 | early | 10.1111/tpj.12723 |
| MtMCA8 | Medtr7g100110 | early | 10.1073/pnas.1107912108 |
| MtNOPE1 | Medtr3g093270 | early | 10.1038/nplants.2017.73 |
| MtPDR1 | Medtr3g097560 | early | 10.1038/nature10873 |
| MtPP2AB’1 | Medtr1g112940 | early | 10.1104/pp.114.246371 |
| MtAnn1 | Medtr8g038210 | early | 10.18388/abp.2009_2451 |
| MtSkl1 | Medtr0041s0030 | early | 10.1111/j.1365-313X.2008.03531.x |
| MtAMT2 | Medtr8g095040 | mature | 10.1105/tpc.114.131144 |
| mtD27 | Medtr1g471050 | mature | 10.1186/s12870-015-0651-x |
| MtDELLA1 | Medtr3g065980 | mature | 10.1073/pnas.1308973110 |
| MtDXS2 | Medtr8g068265 | mature | 10.1111/j.1365-313X.2008.03575.x |
| MtERF1 | Medtr1g078400 | mature | 10.1104/pp.15.01155 |
| MtEXO70i | Medtr1g017910 | mature | 10.1016/j.cub.2015.06.075 |
| MtFatM | Medtr1g109110 | mature | 10.1111/nph.14533 |
| MtGint1 | Medtr1g090920 | mature | 10.1016/j.cub.2021.03.067 |
| MtHA1 | Medtr8g006790 | mature | 10.1105/tpc.113.120436 |
| MtKIN3 | Medtr7g116650 | mature | 10.1111/tpj.15685 |
| MtLEC5 | Medtr5g031030 | mature | 10.1007/s00425-006-0262-8 |
| MtPT4 | Medtr1g028600 | mature | 10.1105/tpc.004861 |
| MtRAD1 | Medtr4g104020 | mature | 10.1104/pp.15.01155 |
| MtRAM1 | Medtr7g027190 | mature | 10.1038/cr.2013.167 |
| MtRAM2 | Medtr1g040500 | mature | 10.1111/nph.14533 |
| MtRFCb | Medtr3g118160 | mature | 10.1038/nplants.2015.208 |
| MtSTR2 | Medtr5g030910 | mature | 10.1105/tpc.110.074955 |
| MtSWEET1b | Medtr3g089125 | mature | 10.1111/nph.15975 |
| MtVAPYRIN | Medtr6g027840 | mature | 10.1111/j.1365-313X.2009.04072.x |
| MtMYB1 | Medtr7g068600 | mature | 10.1016/j.cub.2017.03.003 |
| MtDefMd1 | Medtr8g012805 | late | 10.1371/journal.pone.0191841 |
| MtDefMd3 | Medtr8g012835 | late | 10.1371/journal.pone.0191841 |

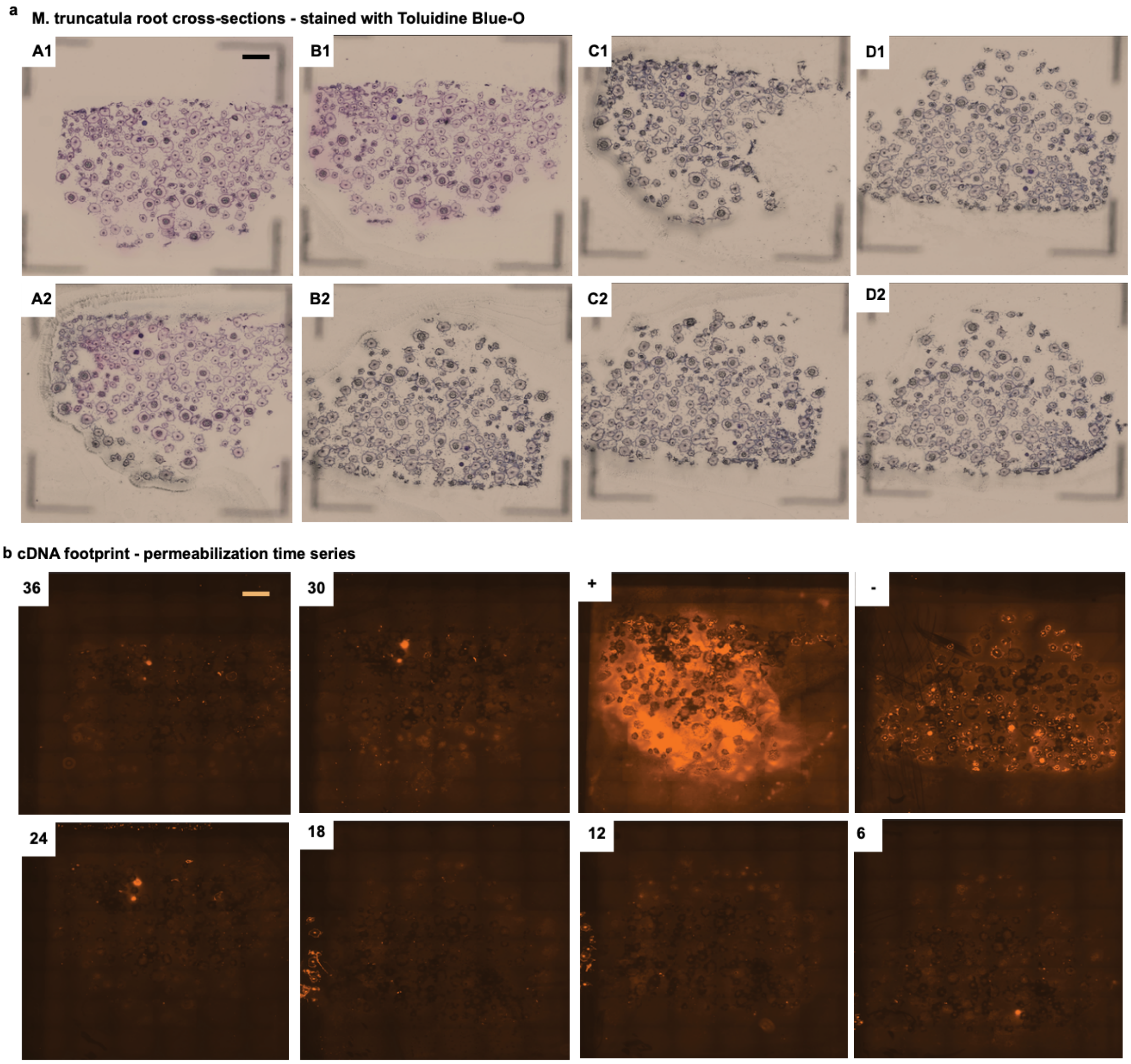


**Supplementary Figure 1: Visium tissue optimization**

**a,** Brightfield images of eight capture areas on final Visium Tissue Optimization slide containing 16 $\mu m$- thick *M. truncatula* lateral root cryo-sections after methanol fixation and staining with Toluidine Blue-O (scale bar = 1mm).
**b,** Corresponding cDNA footprint after fluorescently-tagged nucleotides were incorporated into reverse transcription reaction across 6 different permeabilization timepoints (scale bar = 1mm). Timepoint of 12 minutes was chosen for subsequent Visium Gene Expression workflows.

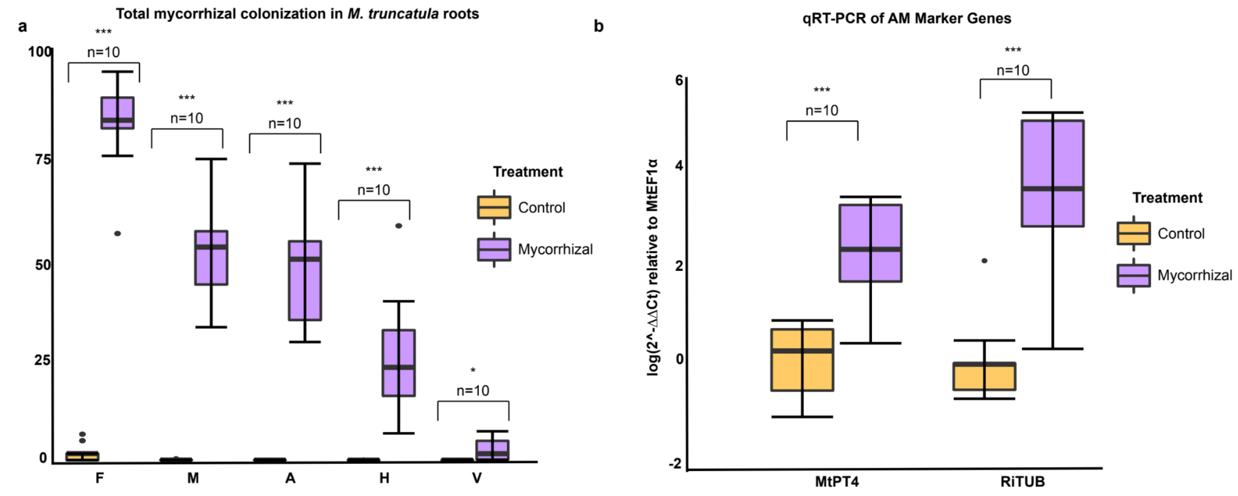


**Supplementary Figure 2: Root colonization analyses**

**a,** AM colonization quantified by the Trouvelot method at 28 dpi with *R. irregularis* in roots; mean ± SEM (n = 10). F%, frequency of infection; M%, total mycorrhization, A%, total arbuscule abundance; H%, total intraradical hyphae abundance, V%, total vesicle abundance.

**b,** Expression of *MtPT4* and RiTUB relative to *MtEF1-a* measured by qRT-PCR; mean ± SEM (n = 10).

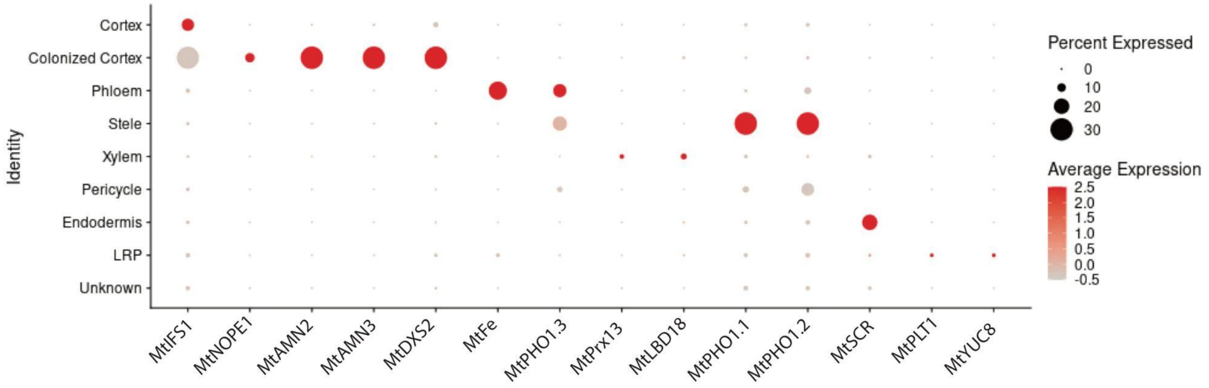


**Supplementary Figure 3: Select marker gene expression for single-nuclei clusters**

Dotplot of expression profiles for select *M. truncatula* marker genes for unique root cell types across labeled cluster identities utilizing hierarchical clustering within the integrated single-nuclei datasets.

**a** **All single-nuclei datasets UMAP - integrated object labeled by cell identity**

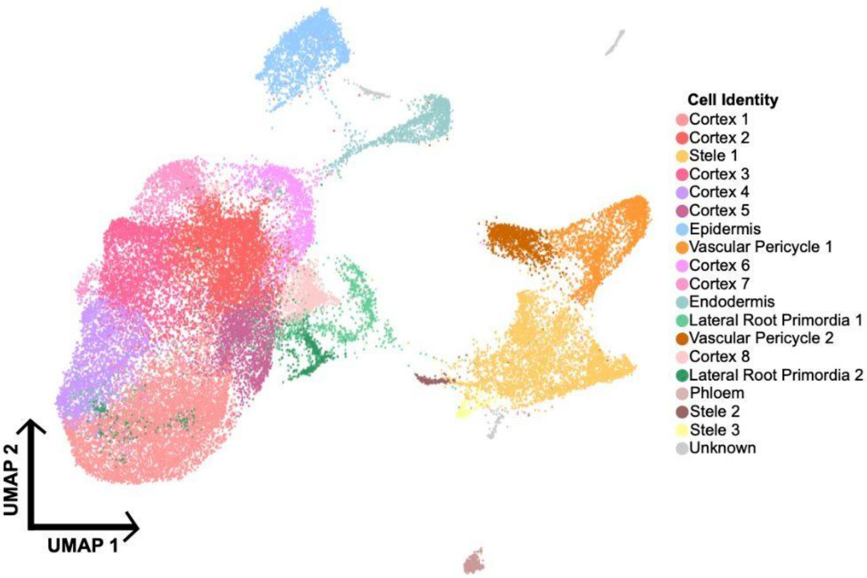


**b** **All single-nuclei datasets UMAP - integrated object labeled by dataset**

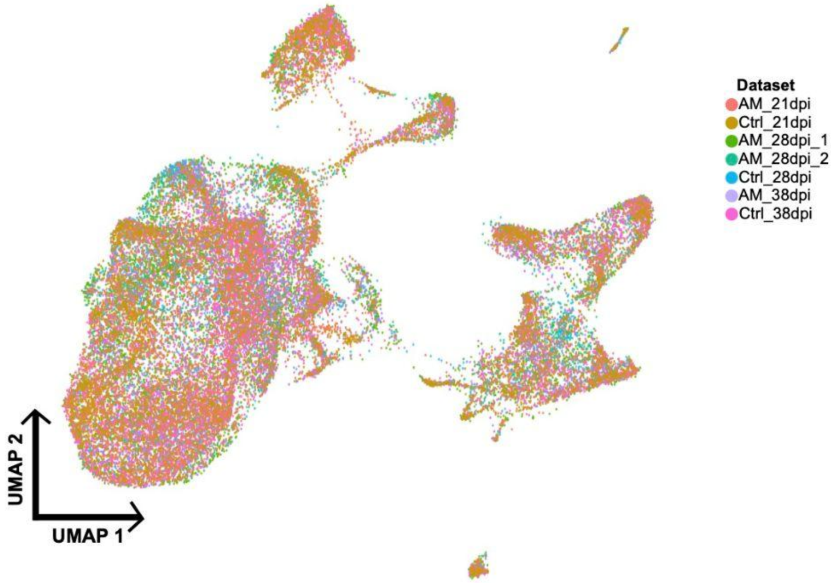


**c**

**Cell number per cluster in each dataset**

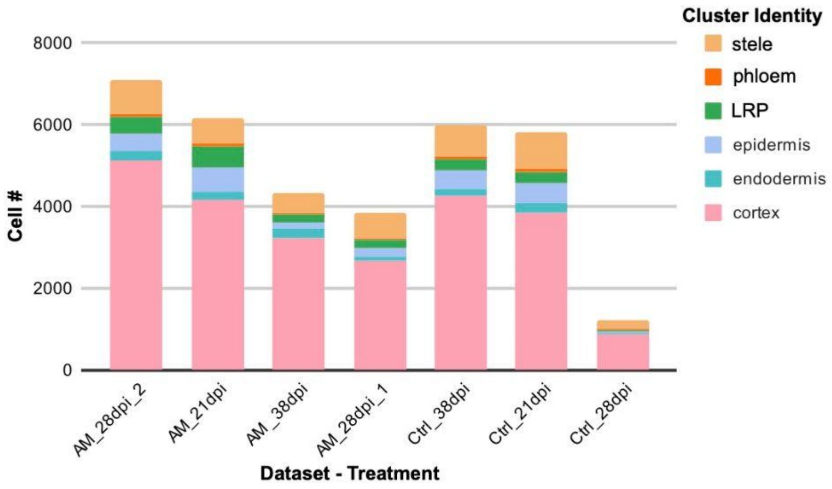

**Supplementary Figure 4: Single-nuclei RNA-seq integrated UMAPs by treatment and dataset**

**a,** UMAP of 38,096 M. *truncatula* nuclei from the integrated Seurat object of all root harvests. The identities of 19 unique clusters are represented by different colors according to cell type. Clusters without a known identity are marked as "unknown".

**b,** UMAP of 38,096 M. *truncatula* nuclei from the integrated Seurat object of all root harvests. Different colors correspond to the original dataset.

**c,** Barplot exhibiting the number of cells that can be assigned to a specific cell type identity for each of the individual datasets within the integrated single nuclei dataset.

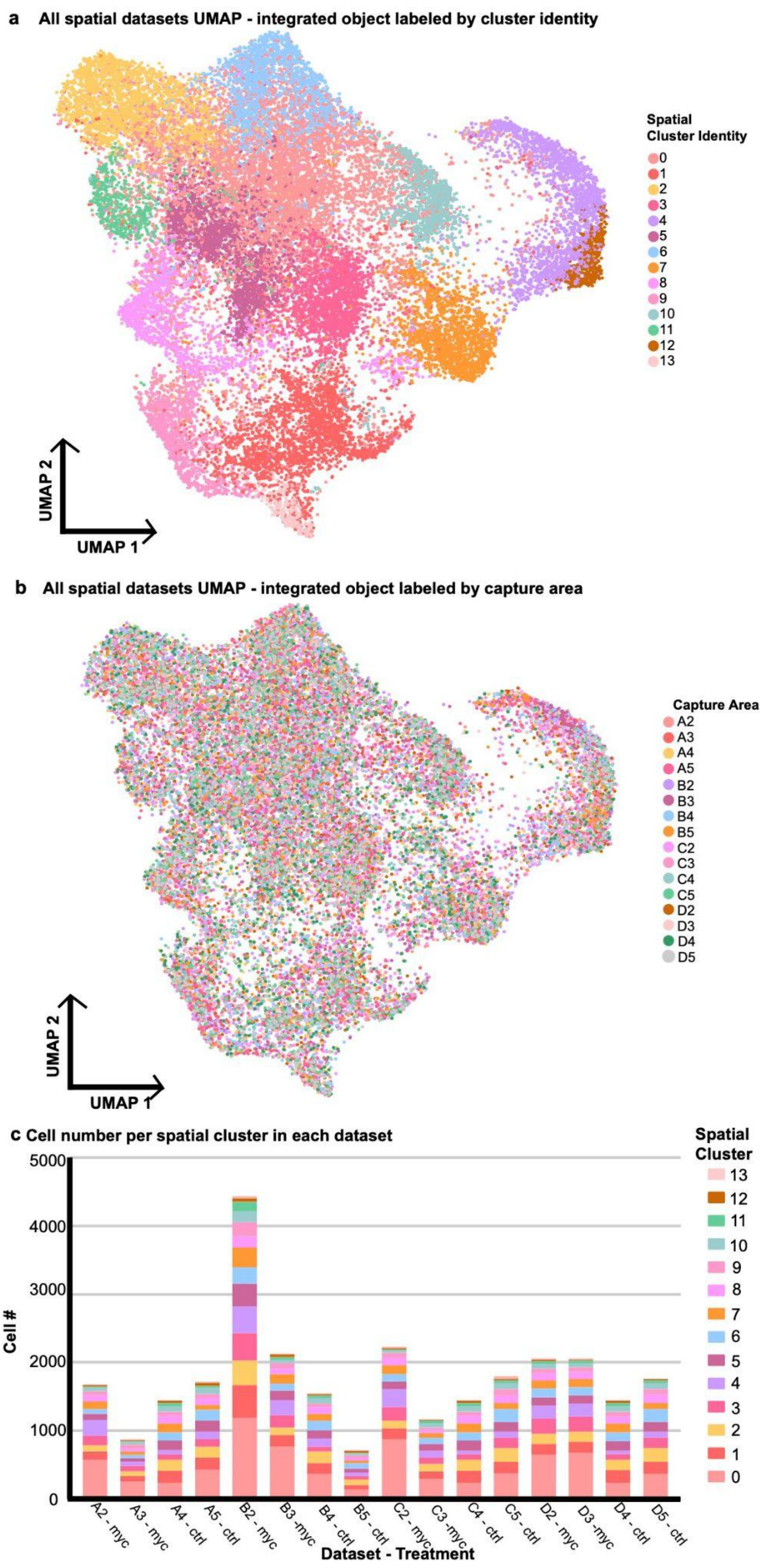
a All spatial datasets UMAP - integrated object labeled by cluster identity
Spatial Cluster Identity
0
1
2
3
4
5
6
7
8
9
10
11
12
13
UMAP 2
UMAP 1
b All spatial datasets UMAP - integrated object labeled by capture area
Capture Area
A2
A3
A4
A5
B2
B3
B4
B5
C2
C3
C4
C5
D2
D3
D4
D5
UMAP 2
UMAP 1
c Cell number per spatial cluster in each dataset
Spatial Cluster
13
12
11
10
9
8
7
6
5
4
3
2
1
0
5000
4000
3000
2000
1000
0
Cell #
A2 - myc
A3 - myc
A4 - ctrl
A5 - ctrl
B2 - myc
B3 -myc
B4 - ctrl
B5 - ctrl
C2 - myc
C3 -myc
C4 - ctrl
C5 - ctrl
D2 - myc
D3 -myc
D4 - ctrl
D5 - ctrl
Dataset - Treatment

**Supplementary Figure 5: Spatial RNA-seq integrated UMAPs by treatment and dataset**
**a,** UMAP of 28,564 spatial voxels from the integrated Seurat object of all root harvests. The identities of 17 unique clusters are represented by different colors according to cluster identity.

**b,** UMAP of 28,564 spatial voxels from the integrated Seurat object of all root harvests. Different colors correspond to the original dataset.

**c,**Barplot exhibiting the number of cells that can be assigned to a specific cluster for each of the individual capture areas within the integrated spatial dataset.

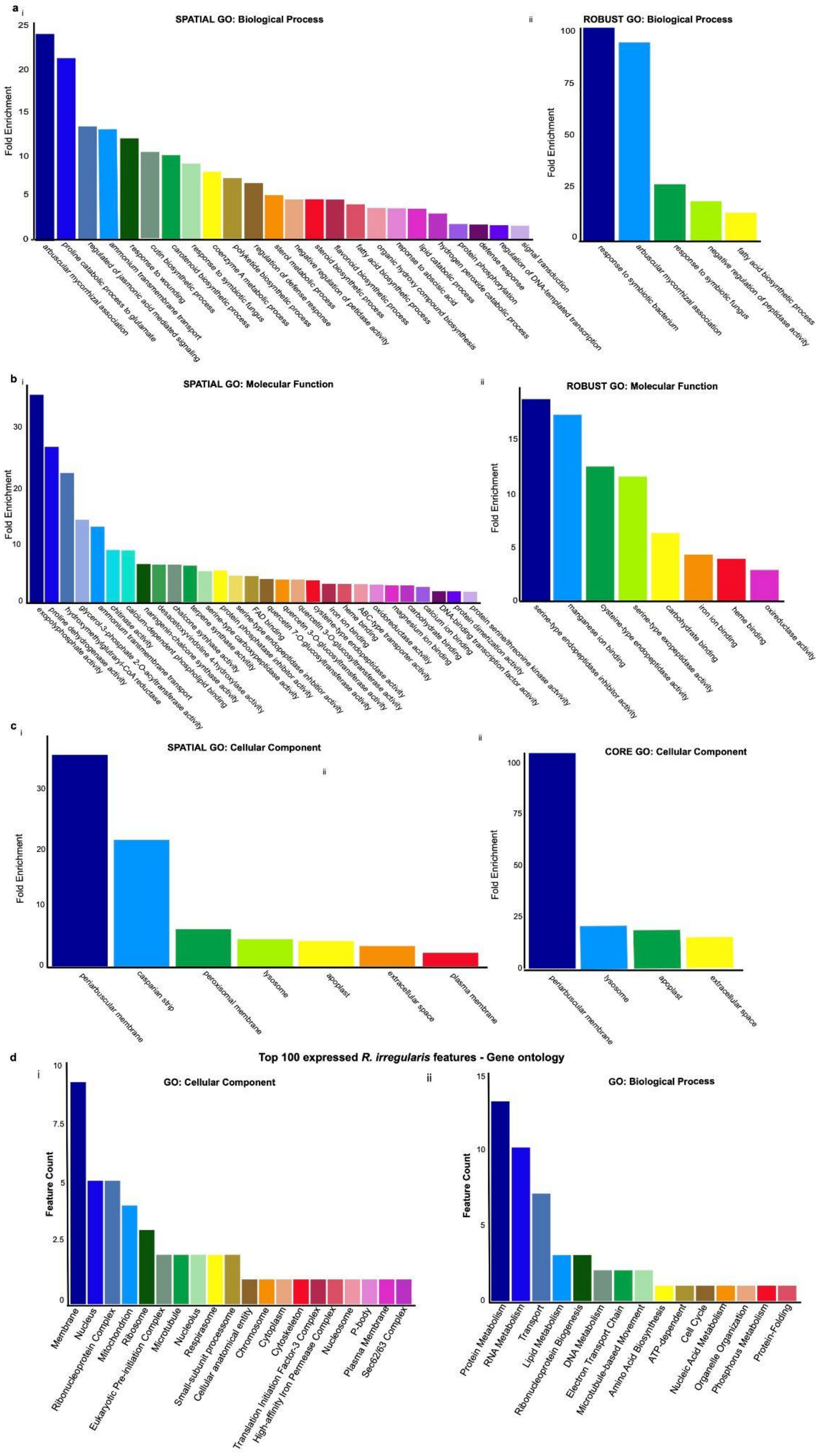
a
i
SPATIAL GO: Biological Process
Fold Enrichment
0
5
10
15
20
25
arbuscular mycorrhizal association
proline catabolic process to glutamate
regulated of jasmonic acid mediated signaling
ammonium transmembrane transport
response to wounding
cutin biosynthetic process
carotenoid biosynthetic process
response to symbiotic fungus
coenzyme A metabolic process
polyketide biosynthetic process
regulation of defense response
sterol metabolic process
negative regulation of petidase activity
steroid biosynthetic process
flavonoid biosynthetic process
fatty acid biosynthetic process
organic hydroxy compound biosynthesis
reponse to abscisic acid
lipid catabolic process
hydrogen peroxide catabolic process
protein phosphorylation
defense response
regulation of DNA-templated transcription
signal transduction
ii
ROBUST GO: Biological Process
Fold Enrichment
0
25
50
75
100
response to symbiotic bacterium
arbuscular mycorrhizal association
response to symbiotic fungus
negative regulation of peptidase activity
fatty acid biosynthetic process
b
i
SPATIAL GO: Molecular Function
Fold Enrichment
0
10
20
30
exopolyphosphate activity
proline dehydrogenase activity
hydroxymethylglutraryl-CoA reductase
glycerol-3-phosphate 2-O-acyltransferase activity
ammonium transmembrane transport
chitinase activity
calcium-dependent phospholipid binding
naringenin-chalcone synthase activity
desacetoxyvindoline 4-hydroxylase activity
chalcone synthase activity
terpene synthase activity
serine-type carboxypeptidase activity
protein phosphatase inhibitor activity
serine-type endopeptidase inhibitor activity
FAD binding
quercetin 7-O-glucosyltransferase activity
quercetin 3-O-glucosyltransferase activity
cysteine-type endopeptidase activity
iron ion binding
heme binding
ABC-type transporter activity
oxidoreductase activity
magnesium ion binding
carbohydrate binding
calcium ion binding
DNA-binding transcription factor activity
protein dimerization activity
protein serine/threonine kinase activity
ii
ROBUST GO: Molecular Function
Fold Enrichment
0
5
10
15
serine-type endopeptidase inhibitor activity
manganese ion binding
cysteine-type endopeptidase activity
serine-type exopeptidase activity
carbohydrate binding
iron ion binding
heme binding
oxireductase activity
c
i
SPATIAL GO: Cellular Component
ii
Fold Enrichment
0
10
20
30
periarbuscular membrane
casparian strip
peroxisomal membrane
lysosome
apoplast
extracellular space
plasma membrane
ii
CORE GO: Cellular Component
Fold Enrichment
0
25
50
75
100
periarbuscular membrane
lysosome
apoplast
extracellular space
d
Top 100 expressed R. irregularis features - Gene ontology
i
GO: Cellular Component
Feature Count
0
2.5
5
7.5
10
Membrane
Nucleus
Ribonucleoprotein Complex
Mitochondrion
Ribosome
Eukaryotic Pre-initiation Complex
Microtubule
Nucleolus
Respirasome
Small-subunit processome
Cellular anatomical entity
Chromosome
Cytoplasm
Cytoskeleton
Translation Initiation Factor-3 Complex
High-affinity Iron Permease Complex
Nucleosome
P-body
Plasma Membrane
Sec62/63 Complex
ii
GO: Biological Process
Feature Count
0
5
10
15
Protein Metabolism
RNA Metabolism
Transport
Lipid Metabolism
Ribonucleoprotein Biogenesis
DNA Metabolism
Electron Transport Chain
Microtubule-based Movement
Amino Acid Biosynthesis
ATP-dependent
Cell Cycle
Nucleic Acid Metabolism
Organelle Organization
Phosphorus Metabolism
Protein-Folding

**Supplementary Figure 6: Gene-set enrichment analysis reveals AM-specific gene functionalities within the mycorrhizal spatial datasets and robust gene list.**

**a,** Gene ontology enrichment scores for the main biological processes represented within the significantly upregulated genes between the mycorrhizal and control treated i) spatial datasets and ii) robust gene list as compared to the reference genome.

**b,** Gene ontology enrichment scores for the main molecular functions represented within the significantly upregulated genes between the mycorrhizal and control treated i) spatial datasets and ii) robust gene list as compared to the reference genome.

**c,** Gene ontology enrichment scores for the main cellular component represented within the significantly upregulated genes between the mycorrhizal and control treated i) spatial datasets and ii) robust gene list as compared to the reference genome.

**d,** Gene ontology categories cellular component (i) and biological process (ii) breakdown identified using the Blast2Go software package for the top 100 most highly expressed *R. irregularis* genes across all nine mycorrhizal capture areas. No annotation could be identified for 44 of the 100 genes.

### Supplementary Text 1

### Cluster Identification for Single-nuclei RNAseq

Arbuscules form in cortical cells, and we determined that clusters 0, 1, 2, 4, 6, 8, 9, 14, and 16 are cortical cells based on enrichment of *ISOFLAVONE SYNTHASE1* (*MtIFS1*, Medtr4g088195) and *ISOFLAVONE SYNTHASE3 (MtIFS3*, Medtr4g088160)[1], as well as *ATP-BINDING CASSETTE TRANSPORTER59* (*MtABCG59*, Medtr3g107870)[2], which encodes a strigolactone transporter which is expressed in cortical cells under phosphate-depleted conditions. Cluster 14 represents cortical cells which are colonized by AM fungi, based on a range of known marker genes, including *ATP-BINDING CASSETTE TRANSPORTER* (*MtNOPE1*, Medtr3g093270)[3], two isoforms of *ABCB FOR MYCORRHIZATION AND NODULATION* (*MtAMN2,* Medtr4g081190 and *MtAMN3,* Medtr8g022270)[4], *1-deoxy-D-xylulose 5-phosphate synthase* (*MtDXS2,* Medtr8g068265)[5], and *REDUCED ARBUSCULAR MYCORRHIZA1 (MtRAM1,* Medtr7g027190[6]), and *VAPYRIN* (*MtVPY,* Medtr1g089180)[7]. We specifically focused on more mature roots, excluding meristematic or developing cells for most cell types as arbuscules do not form in these cell types. As a result, we did not observe evidence for developmental variation (trajectories) that are typically captured in single-cell studies of embryonic or meristematic tissues. Marker genes for quiescent center and lateral root primordia, however, were used to identify cluster 12 as meristematic cells, including four homologs of *PLETHORA* (*MtPLT1-4,* Medtr2g098180, Medtr4g065370, Medtr5g031880, and Medtr7g080460)[8] and *YUCCA (MtYUC8,* Medtr7g099330)[9] . *MtYUC8* and *MtPLT* genes tend to be associated with nodule formation, as well, but other genes which are upregulated by nodulation, such as NODULE INCEPTION (*MtNIN,* Medtr5g099060)[9], *MtRPG*[10] (Medtr1g090807), FLOTILLIN-LIKE (*MtFLOT,* Medtr3g106430)[11] were either absent from our dataset or expressed at low levels and not specific to any cluster. RESPIRATORY BURST OXIDASE HOMOLOGS (*MtRboHF,* Medtr7g060540)[12], which is specifically expressed in root hairs, defined a small cluster adjacent to the LRP cluster as root hairs. The presence of *SCARECROW* (*MtSCR*, Medtr7g074650) indicated that cluster 7 represents endodermal cells[13]. Clusters 3, 5, and 11 were predicted to be vascular tissue, with several stele-specific arabidopsis homologs such as three homologs of transcription factor MYB DOMAIN PROTEIN (MtMYB0*71*, Medtr5g014990, *MtMYB113* Medtr2g096380, and *MtMYB112*, Medtr4g063100)[14], as well as functionally characterized *M. truncatula* marker genes enriched in these clusters: three PHOSPHATE TRANSPORTER homologs: (*MtPHO1.1-1.3,* Medt1g041695, Medtr1g075640, Medtr8g069955)[15] and *SUGAR TRANSPORT PROTEIN13* (*MtSTP13,* Medtr1g104780)[16]. Based on homologs from Arabidopsis marker genes such as PEROXIDASE13 (*MtPrx13*, Medtr1g101830) and LOB-DOMAIN PROTEIN (*MtLBD18,* Medtr8g036085)[14] , cluster 17 represents xylem cells. *FE/ALTERED PHLOEM DEVELOPMENT* (*MtFe,* Medtr6g444980) and Arabidopsis PHLOEM EARLY DOF1 homologs (*PEAR1*, Medtr3g077750 and Medtr4g461080)[17] were enriched in cluster 13, suggesting that these are phloem cells. We defined cluster 15 as representing companion cells, as it is enriched for Arabidopsis phloem marker homolog SUPER NUMERIC NODULES (*MtSUNN,* Medtr4g070970)[18] and homologs of Arabidopsis companion cell markers Arabidopsis *SUCROSE PROTON SYMPORTER2* (*AtSUC2)* (Medtr1g096910) and Arabidopsis *FT INTERACTING PROTEIN1 (AtFTIP1,* Medtr0291s0010)[19]. Clusters 5 and 11 were enriched for *MtPHO1.1-1.3* (Medtr1g041695, Medtr1g075640, and Medtr8g069955), suggesting that these are central cylinder/pericycle cells[14,15].
